# Deep sequencing of nonenzymatic RNA primer extension

**DOI:** 10.1101/2020.02.18.955120

**Authors:** Daniel Duzdevich, Christopher E. Carr, Jack W. Szostak

**Affiliations:** Department of Molecular Biology, Center for Computational and Integrative Biology, Massachusetts General Hospital, Boston, MA 02114, United States of America; Howard Hughes Medical Institute, Massachusetts General Hospital, Boston, MA 02114, United States of America; Department of Earth, Atmospheric and Planetary Sciences, Massachusetts Institute of Technology, Cambridge, MA, 02139, United States of America; Department of Chemistry and Chemical Biology, Harvard University, Cambridge, MA 02138, United States of America; Department of Genetics, Harvard Medical School, Boston, MA 02115, United States of America

## Abstract

Life emerging in an RNA world is expected to propagate RNA as hereditary information, requiring some form of primitive replication without enzymes. Nonenzymatic template-directed RNA primer extension is a model of the polymerisation step in this posited form of replication. The sequence space accessed by primer extension dictates potential pathways to self-replication and, eventually, ribozymes. Which sequences can be accessed? What is the fidelity of the reaction? Does the recently-illuminated mechanism of primer extension affect the distribution of sequences that can be copied? How do sequence features respond to experimental conditions and prebiotically relevant contexts? To help answer these and related questions, we here introduce a deep-sequencing methodology for studying RNA primer extension. We have designed and vetted special RNA constructs for this purpose, honed a protocol for sample preparation and developed custom software that sorts and analyses raw sequencing data. We apply this new methodology to proof-of-concept controls, and demonstrate that it works as expected and reports on key features of the sequences accessed by primer extension.

## INTRODUCTION

A central challenge of the RNA world hypothesis is replicating RNA without enzymes (1–7). Replication, in turn, requires a copying mechanism. NonEnzymatic RNA Primer Extension (NERPE) is a model system of template-directed chemical RNA copying in which a primer is extended by the polymerisation of nucleotides or the ligation of oligonucleotides (oligos) (8–10). Standard nucleotides with triphosphate activation do not react spontaneously with a primer-template complex because they are effectively inert without highly evolved enzymes (11). Other activating moieties have been identified (12), including 2-aminoimidazole (2AI) (13, 14). The mechanism of primer extension with imidazole-based activation was only recently illuminated (15–19) (Figure S1A-D). Briefly, monoribonucleotides are chemically activated with 2AI to yield 2AI-monoribonucleotides (2AIrN). In aqueous solution buffered to a pH = pK_a_ of 2AI (= 8.3), 2AIrN react spontaneously to form a highly reactive 5’-5’phospho-imidazolium-phospho bridged dinucleotide intermediate. The dinucleotide intermediate binds the template through Watson-Crick base pairing, and the deprotonated oxygen of the primer 3’ hydroxyl attacks the proximal bridging phosphate displacing a 2AIrN as the leaving group. Interestingly, although the reactive intermediate is a dinucleotide, primer extension proceeds one nucleotide at a time (Figure S1C-D). A complementary pathway is nonenzymatic ligation of an activated oligo, a reaction that proceeds without a bridged intermediate: the deprotonated oxygen of the primer 3’ hydroxyl attacks the activated phosphate, and the activating moiety (for example, 2AI) is the leaving group, yielding a primer extended by the length of the reacting oligo. Both pathways yield canonical 3’-5’ phosphodiester backbone linkages, but 2’-5’ linkages can also form, especially in the context of mismatched bases (20–23). The prebiotic relevance of primer extension as a model of RNA copying assumes that there are effective chemical pathwyas to nucleotides, nucleotide activation chemistry, templates and primers—all topics of ongoing research (24–29).

The prevailing technique for characterising nonenzymatic RNA primer extension is denaturing PolyAcrylamide Gel Electrophoresis (PAGE) (Figure S1E). The primer is fluorescently labelled, and the reaction products (primers extended to different lengths) are separated by size, typically yielding a characteristic banding pattern that can be directly mapped to +1, +2, +3, *etc.* extension events. PAGE analysis can be used to measure reaction kinetics (16), test novel reaction chemistries (14) and help determine mechanisms (19). A significant limitation of PAGE analysis is that information about the sequences accessed by primer extension is strictly determined by defined template and reactant identities. For example, interpreting a hypothetical +1 product as an added rG nucleotide strictly requires rC in the template and activated 2AIrG as the reactant. If the set of available template sequences harbours other bases in that position, and/or if additional activated nucleotides are used as reactants, then PAGE analysis cannot definitively establish the identity of the +1 product. Furthermore, mixtures of activated nucleotides can yield products with mismatches to the templating sequence (errors), and mismatches are difficult or impossible to identify by PAGE analysis. The nature of copying fidelity in primer extension is of great interest (30–33) because there is no protein machinery to control and proofread polymerisation. An ideal deep-sequencing methodology should be applicable to any experimentally-defined templating sequence, should access mismatch information (including relevant sequence context) and should tolerate a wide range of reaction conditions.

Here we introduce NERPE-Seq, a deep-sequencing protocol and analysis pipeline for studying nonenzymatic RNA primer extension. We developed special RNA constructs for this purpose, and a protocol to prepare multiple experimental samples for simultaneous deep-sequencing (multiplexing). We have identified potential sources of bias arising from enzymatic reactions, RNA backbone heterogeneity and RNA secondary structure. We then systematically addressed these biases to ensure that sequencing data reflect the properties of primer extension rather than any incidental selectivities of the protocol. We also created custom analysis software that filters and sorts raw data, and generates standard measurements of template and product sequences. We used a set of control constructs and reactions to demonstrate that NERPE-Seq accurately reproduces known results, especially the extent of primer extension as measured by PAGE analysis, the currently accepted standard. We show that NERPE-Seq can characterise both nonenzymatic polymerisation and ligation reactions, and identify mismatches. Finally, we assess the potential sources and magnitude of noise in the data. We conclude with a summary of our validations and emphasise potential applications for studying nonenzymatic copying in the context of the RNA world hypothesis.

## MATERIAL AND METHODS

### General

All reactions were performed with RNase-free water and salt solutions (not DEPC-treated, Ambion™) in RNase-free 0.2 ml PCR tubes (VWR International) or 1.5 ml DNA LoBind tubes (Eppendorf). Bicine buffer was prepared from the Na^+^ salt (Sigma-Aldrich) with RNase-free UltraPure™ Distilled Water (Invitrogen™), adjusted to pH = 8 with 2 M NaOH and syringe-filtered (Millex^®^ MP 0.22 μm, Millipore Sigma). Extreme caution was taken to prohibit contamination, including the use of barrier tips for all liquid handling (MultiMax, Sorenson™ BioScience, or TipOne^®^, USA Scientific) except non-preparative gel loading. Enzymes and DNA/RNA ladders were purchased from New England BioLabs^®^ (NEB), and used as per the manufacturer’s instructions unless otherwise indicated. All incubations with an indicated temperature were performed in a Bio-Rad T100™ thermal cycler.

### Synthesis of 2-aminoimidazole-activated monoribonucleotides

0.63 mmole (1 equivalent) of the nucleoside-5’-monophosphate free acid (Santa Cruz Biotechnology or ACROS Organics) and 5 equivalents of 2-aminoimidazole hydrochloride (Combi-Blocks) were dissolved in 30 ml UltraPure™ Distilled Water and the pH adjusted to 5.5 with syringe-filtered 1M NaOH. The mixture was syringe-filtered and aliquoted into three 50 ml polypropylene Falcon tubes (Corning), flash frozen in liquid nitrogen and lyophilised. 30 ml dry dimethyl sulfoxide (DMSO, Sigma-Aldrich) and 1.2 ml dry triethylamine (TEA, Sigma-Aldrich) were mixed in a dry 250 ml glass roundbottom flask with a stir-bar (baked for at least an hour at 120°C and then cooled in a desiccator with Drierite) under argon; the lyophilised nucleoside-5’-monophosphate and 2-aminoimidazole hydrochloride were added, and the mixture was sonicated and gently heated until it turned clear. To this was added 9 equivalents of triphenylphosphine (Sigma-Aldrich), completely dissolved by sonication, and then 10 equivalents of 2,2’-dipyridyldisulfide (DPDS, Combi-Blocks); the mixture was stirred under argon for 30 minutes. The mixture was poured into a ice-chilled solution of 400 ml acetone (Fisher Scientific), 250 ml diethyl ether (Fisher Scientific), 30 ml TEA and 1.6 ml acetone saturated with NaClO_4_ (Sigma-Aldrich) in a glass bottle with stirring. The target material quickly flocculated. The stirring was stopped and the bottle put back on ice for approximately 30 minutes. Most of the supernatant was removed with a glass pipette attached to a vacuum trap, leaving slightly less than 200 ml. The remaining mixture with the flocculant was aliquoted into four Falcon tubes and centrifuged at 2,500 rpm for 3 minutes (the centrifugation parameters are not critical so long as the supernatant clarifies and the pellet is compact enough to withstand decanting). The supernatant was discarded, and the pellet resuspended by vortexing (washed) in a solution of 1:8.3:13.3 of TEA:diethyl ether:acetone and centrifuged again. The wash was repeated twice with just acetone; the final wash was performed in two 50 ml Bio-Reaction tubes (Celltreat^®^) with gas-permeable membranes in the caps, and the pellets dried overnight under vacuum. Each sample was dissolved in 5 ml UltraPure™ Distilled Water, and the combined volume was purified by reverse-phase flash chromatography (CombiFlash^®^, Teledyne ISCO) over a 50 g RediSep Rf Gold C18Aq column (Teledyne ISCO) at 40 ml/min. using gradient elution between (A) Milli-Q^®^ water (Milli-Q^®^ Reference with Biopak^®^ Polisher, Millipore Sigma) and (B) acetonitrile (Fischer Scientific). Absorbance was monitored at 214 nm and 260 nm. The target activated monoribonucleotide eluted as a dominant and baseline-separated peak at approximately five column volumes of a ten column volume elution between 0% and 10% (B). Target fractions were quickly placed on ice, pooled, adjusted to pH = 10 with 1 M NaOH, flash frozen in Falcon tubes with no more than 20 ml per tube, lyophilised and stored at -80°C. Aliquots for experiments were prepared by resuspending a lyophilised sample in a small volume of water (typically less than 500 μl), syringe-filtering, using 2 μl to prepare a 1:100 dilution and then measuring the absorbance at 260 nm with a NanoDrop™ 2000c spectrophotometer (ThermoFisher Scientific). Further dilutions in 1:10 increments were used to ensure a linear range measurement. Established absorbances (34) were used to calculate the stock concentration, and the stock was aliquoted into 1.5 ml tubes at a volume equivalent to 0.002 mmole. The aliquots were frozen on dry ice, lyophilised at -20°C and stored at -80°C. Product identity and purity were confirmed by ^31^P NMR on a 400 MHz spectrometer operating at 161 mHz (Varian INOVA). Spectra were analysed by MNova 11 (Mestrelab). Lyophilised samples were resuspended in 10 μl water immediately prior to use.

### Synthesis of 2-aminoimidazole-activated 5’-CGG-3’ (2AI-CGG) trimer

The 5’-phosphorylated trimer was synthesised at the 50 μmole scale on a universal support (Glen Research) using standard RNA phosphoramidites (BioAutomation) and diethoxy N,N-diisopropyl phosphoramidite phosphorylating reagent (ChemGenes) on a MerMade synthesiser (BioAutomation). The product was cleaved from the support with 15 ml of a 1:1 mixture of 28-30% ammonium hydroxide (Sigma-Aldrich) and 40% aqueous methylamine (Sigma-Aldrich), and the mixture split into three 15 ml Falcon tubes. The oligo was deprotected by incubation at 65°C for 2.5 hours, then vacuum centrifuge concentrated (SpeedVac, ThermoFisher Scientific) for two hours until 20-30% of the initial volume was left. The samples were flash frozen and lyophilised overnight. The lyophilised samples were each resuspended in 5 ml of a 1:1 mixture of DMSO:TEA/3HF (Sigma-Aldrich), incubated at 65°C for 2.5 hours, allowed to come to room temperature and pooled in a 50 ml Falcon tube. The oligo was precipitated with 625 μl of 3 M sodium acetate (Sigma-Aldrich) and 15 ml 1-butanol (Sigma-Aldrich). The mixture was centrifuged at 4,000 rpm for 5 minutes, the supernatant discarded, and the pellet washed twice with ethanol and dried overnight under vacuum.

Each sample was dissolved in 5 ml of 25 mM triethylamine bicarbonate, pH = 7.5 (1M stock prepared by bubbling CO_2_ into a solution of Milli-Q^®^ water and TEA), and purified by reverse-phase flash chromatography over a 50 g RediSep Rf Gold C18Aq column at 40 ml/min. using gradient elution between (A) 25 mM TEAB, pH = 7.5 and (B) acetonitrile. Absorbance was monitored at 214 nm and 260 nm. The target trimer eluted as a dominant and baseline-separated peak at approximately five column volumes of a ten column volume elution between 0% and 20% (B). Target fractions were pooled and concentrated to 3-5 ml in a 250 ml flask on a rotary evaporator with a water bath set to 32°C. The material was transferred to a 50 ml Falcon tube, flash frozen and lyophilised overnight.

The purified trimers were activated using the same approach as for monoribonucleotides above with minor adjustments. ∼10 μmole of the trimer was resuspended in 5 ml dry DMSO and 56 μl dry TEA in a dry 50 ml glass roundbottom flask with a stir-bar. To this was added 30 equivalents each of 2-aminoimidazole hydrochloride, triphenylphosphine and DPDS; the mixture was stirred under argon for five hours. The mixture was poured into an ice-chilled solution of 80 ml acetone, 50 ml diethyl ether, 6 ml TEA and 0.32 ml acetone saturated with NaClO_4_ in a glass bottle with stirring. The target material quickly flocculated. The stirring was stopped and the bottle put back on ice for approximately 30 minutes. Most of the supernatant was removed with a glass pipette attached to a vacuum trap, leaving slightly less than 40 ml. The remaining mixture with the flocculant was aliquoted into four Falcon tubes and centrifuged at 3,000 rpm for 3 minutes. The supernatant was discarded, and the pellet resuspended by vortexing in a solution of 1:9.3:22.6 of TEA:diethyl ether:acetone and centrifuged again. The wash was repeated twice with just acetone; the final wash was performed in two 50 ml Bio-Reaction tubes and the pellets dried overnight under vacuum. The activated trimer was purified by HPLC (Agilent) over an Eclipse XDB C18 column (Agilent) at 10 ml/min. using gradient elution between (A) 25 mM TEAB, pH = 7.5 and (B) acetonitrile. Absorbance was monitored at 260 nm. The target activated trimer eluted as a dominant and baseline-separated peak during a 20 minute elution between 3% and 12% (B). Target fractions were pooled and the concentration measured with a NanoDrop™ spectrophotometer. The stock was aliquoted into 1.5 ml tubes at a volume equivalent to 0.032 μmole, flash frozen, lyophilised overnight at -20°C and stored at -80°C. Product identity and purity were confirmed by Time-Of-Flight Liquid Chromatography-coupled Mass Spectroscopy (TOF LC-MS, Agilent 6230). Lyophilised samples were resuspended in 32 μl water immediately prior to use.

### Oligonucleotides

Oligonucleotides (oligos) were ordered from Integrated DNA Technologies (IDT), NEB or synthesised in-house. All oligos were stored in TE (10 mM Tris-Cl, 1 mM EDTA) buffer, pH = 7 (Invitrogen™) and at -30°C. A table of all oligos used in this study is included in the Supplementary data (Table of Oligonucleotides).

Oligos synthesised in-house, excepting the trimer described above, were prepared on an ABI Expedite instrument (standard phosphoramidites from ChemGenes; special phosphoramidites, synthesis reagents and supports from Glen Research) at the 1 μmole scale with DMT-on for purification. The product was cleaved from the support with 1.3 ml of a 1:1 mixture of 28-30% ammonium hydroxide and 40% aqueous methylamine. The bases were deprotected by incubation at 65°C for 10 minutes, then vacuum centrifuge concentrated for two hours until ∼50% of the initial volume was left. The sample was flash frozen and lyophilised overnight at -20°C. The lyophilised material was resuspended in 115 μl DMSO, 60 μl TEA and 75 μl TEA/3HF, incubated at 65°C for 2.5 hours, and allowed to come to room temperature. The oligo was then purified with a GlenPak (Glen Research) using Norm-Ject^®^ syringes (Air-Tite), eluted with 200 mM ammonium bicarbonate (Sigma-Aldrich, from a syringe-filtered 1 M stock in UltraPure™ Distilled Water) + 30% v/v acetonitrile (Fisher Scientific) (the first one or two drops and the final foam were discarded), vacuum centrifuge concentrated for 45 minutes, flash frozen in liquid nitrogen and lyophilised overnight at -20°C.

The lyophilised material was resuspended in 500 μl TE, pH = 7 and PAGE-purified. PAGE gels were cast as described below for analytical gels, but with 1.5 mm thick spacers and larger wells. 50 μl of the oligo stock was mixed with 50 μl Urea Load Buffer (8.3 M urea [Sigma-Aldrich], 1.3X TBE buffer [from a 10X autoclaved stock], 75 μM bromophenol blue [Sigma-Aldrich, from a 7.5 mM stock in DMSO], 880 μM orange G [Sigma-Aldrich, from an 88 mM stock in DMSO], syringe-filtered) and subjected to PAGE at 10 W for 15 minutes, then 25 W for ∼1 hour (or longer, depending on the size of the target). One of the two glass plates was removed (leaving the other for the gel to rest on), and 10 ml 1X TBE + 1 μl SYBR™ Gold Nucleic Acid Gel Stain (Invitrogen™) was poured on top of the gel and allowed to incubate for 3-4 minutes before being rinsed off with Milli-Q^®^ water. The gel was imaged with a blue light transilluminator (Safe Imager™ 2.0, ThermoFisher Scientific): the target material typically appeared as unstained due to its extremely high concentration relative to well-stained higher (not fully deprotected) and lower (truncated) molecular weight side-products. The target band was excised with a clean razor, purified with a ZR small-RNA PAGE Recovery Kit (Zymo Research), eluted in 30 μl TE, pH = 7 and quantified with a NanoDrop™ spectrophotometer. Product identity was confirmed by Time-Of-Flight Liquid Chromatography-coupled Mass Spectroscopy (TOF LC-MS, Agilent 6230), except in the case of hairpin constructs, which were too large for accurate mass spectroscopy analysis.

### Standard primer extension reactions

Unless otherwise indicated, 1 μM primer and 1.2 μM template were mixed in an appropriate volume of water and 200 mM Na^+^ bicine, pH = 8, incubated at 85°C for 30 s, then cooled to 23°C at 0.2 °C/s. Activated nucleotides or trimer were added to indicated concentrations (in cases with multiple nucleotide species, they were first mixed separately before addition), and the reaction was initiated by the addition of 50 mM MgCl_2_ (all concentrations indicate final concentrations, typically in a 20 μl volume). Timepoints were quenched by adding 1 μl of the reaction mixture to 20 μl Urea Load Buffer. 2.5 μl of the quenched material was then added to 1 μl of a 300 μM stock of RNA complementary to the template and incubated at 95°C for 3 minutes, then cooled to 25°C at 0.2 °C/s. 16.5 μl additional Urea Load Buffer was added and the sample subjected to PAGE at 5 W for 10 minutes then 15 W for 1 hour. In experiments with NPOM-caged bases in the template, the quench was into dye-free Urea Load Buffer, allowing for UV un-caging prior to the addition of RNA complementary to the template.

### Gels and gel analysis

Polyacrylamide gels were prepared with the SequaGel - Urea Gel system (National Diagnostics). A 50 ml mix to yield a 20% gel was degassed under house vacuum with stirring for ∼10 minutes before the addition of 10 μl tetramethylethylenediamine (TEMED, Sigma-Aldrich) and 100 μl of fresh 10% w/v ammonium persulfate (APS, Sigma-Aldrich) prepared with UltraPure™ Distilled Water. The gel was cast as 0.75 mm thick (1.5 mm for preparative) using 20 x 20 cm plates and allowed to set for two hours. Gels were pre-run for at least 45 minutes at 20 W; all runs were at constant power, with heat-distributive aluminum or iron face-plates, in 1X TBE prepared in Milli-Q^®^ water. Gels requiring staining (in particular, gels used to assay the sequencing constructs, which have no attached dyes) were removed from the glass plates and incubated in ∼140 ml of 1X TBE + 14 μl SYBR™ Gold Nucleic Acid Gel Stain for several minutes. All gels were imaged with a Typhoon 9410 scanner (GE Healthcare) using the Auto PMT setting and at 50 μm resolution. For analysis and visualisation, TIFF-formatted images from the Typhoon were imported into Fiji (35). Bands were quantified using the Gel Analysis function, with relative band intensities reported as the ratio of the band intensity to the total lane intensity. Band edges were excluded. Contrast and color changes were applied uniformly to the entire gel image in all cases.

Non-preparative agarose gels were prepared as 1.4% w/v agarose (UltraPure™ Agarose, Invitrogen™) in 50 ml 1X TAE prepared in Milli-Q^®^ water + 5 μl SYBR™ Safe DNA Gel Stain (Invitrogen™, added after microwave heating to form the agarose solution) and cast as 10 x 5 cm. Samples were loaded with Purple Loading Dye (NEB) and run at 80 V for 55 minutes at constant voltage. Gels were imaged on a ChemiDoc imaging system (Bio-Rad). For visualisation, TIFF-formatted images were imported into Fiji. Contrast and color changes were applied uniformly to the entire gel image in all cases.

### RNA sample preparation for deep-sequencing

An appropriate volume of water, 200 mM Na^+^ bicine, pH = 8, 1 μM hairpin construct and (where indicated) 1.2 μM 5’ Handle Block were incubated at 95°C for 3 minutes and cooled to 23°C at 0.2 °C/s. The activated monoribonucleotides or activated trimer and 50 mM MgCl_2_ were added and the mixture was briefly vortexed. The final volume was 30 μl. The mixture was incubated at 23°C with a 42°C heated lid for 24 hours. The reaction was quenched by the addition of 20 μl water followed by desalting in a MicroSpin G-25 spin column (GE Healthcare). The ∼53 μl eluate was transferred to a PCR tube and placed under a 385 nm UV lamp (∼3 cm distance from the tops of the tubes, Spectroline ENF-240C, Spectronics^®^) for 45 minutes with a manual mixing step halfway. An equal volume of Urea Load Buffer was added, the mixture was heated to 95°C for 3 minutes, cooled to 35°C and loaded on a preparative 20% polyacrylamide gel to completely remove residual unreacted nucleotides or oligos, which can interfere with downstream steps. The sample was subjected to PAGE at 10 W for 15 minutes and then 25 W for 1 hour. The gel was stained in 120 ml TBE with 12 μl SYBR™ Gold Nucleic Acid Gel Stain for 10 minutes, imaged with a blue light transilluminator and the target band was excised with a clean razor, purified with a ZR small-RNA PAGE Recovery Kit, and eluted in 7 μl Tris-Cl, pH = 7. The eluate was mixed with 0.36 μl RT Handle (from a 100 μM stock), heated to 95°C for 3 minutes and cooled to 25°C at 1 °C/s. 1.8 μl 10X RNA Reaction Buffer, 7.2 μl 50% PEG_8000_ and 3 μl T4 RNA Ligase 2, truncated KQ were added and the mixture incubated for 18 hours at 25°C and 2 hours at 4°C with a 42°C heated lid. 0.7 μl 5 M NaCl, 0.5 μl SDS (from a 10% w/v stock) and 0.25 μl Proteinase K were added and the mixture incubated for 20 minutes at room temperature. 50 μl of 250 mM NaCl was added and the solution was extracted with 160 μl phenol-chloroform (UltraPure™ phenol:chloroform:isoamyl alcohol 25:24:1, Invitrogen™) and twice with 200 μl chloroform (EMSURE, Millipore Sigma). The final extracted volume was ∼66 μl, which was run through an Oligo Clean & Concentrator spin column (Zymo Research) and eluted in 12 μl water.

### Preparing and purifying cDNA for deep-sequencing

0.7 μl RT Primer (from a 100 μM stock) was added to the purified sample and incubated at 75°C for 3 minutes, 37°C for 10 minutes and 25°C for 10 minutes. 4 μl of ProtoScript^®^ II Buffer, 1 μl dNTP mix (NEB), 0.2 μl MgCl_2_, 2 μl DTT (NEB) and 2 μl of ProtoScript^®^ II were added and the mixture incubated at 42°C for 12 hours with a 105°C heated lid. (Alternatively, for standard conditions [see section on 2’-5’ linkages below], no MgCl_2_ was added and the incubation was at 50°C for 1 hour.) 10 μl of water was added and the mixture was run through an Oligo Clean & Concentrator spin column, including the RNA degradation step as per the manufacturer’s instructions. The cDNA was eluted in 30 μl of TE, pH = 7 and stored at 4°C. The cDNA stock concentration was measured with a NanoDrop™ spectrophotometer, and typically found to be in the fraction of a μg/μl range.

### Preparation and isolation of barcoded PCR products for multiplexed deep-sequencing

A volume of isolated cDNA sample equivalent to 0.1 μg total cDNA was added to a 50 μl Q5^®^ Hot Start High-Fidelity DNA Polymerase PCR reaction with 0.2 μM each of the NEBNext^®^ SR Primer for Illumina^®^ and NEBNext^®^ Index Primer for Illumina^®^ and run for 6 total cycles with a 15 s and 62°C extension step. 10 μl Purple Loading Dye was added to the PCR product mixture and the entire volume was run on a preparative agarose gel. Gels were prepared as 1.4% w/v agarose (Certified Molecular Biology Grade, Bio-Rad) in 1X TAE (UltraPure™, Invitrogen™) and 4 μl SYBR™ Safe DNA Gel Stain, and cast as 10 x 5 cm with large wells sufficient to hold the 60 μl final sample volume. The gel was run at 70 V for 90 minutes at constant voltage. Target bands were excised on a blue light transilluminator and purified with a Quantum Prep^®^ Freeze ‘N Squeeze spin column (Bio-Rad). The eluate was further purified with a magnetic bead clean-up (Agencourt^®^ AMPure^®^ XP) with a 1.7:1 ratio of beads to sample volume. Target material was eluted with 20 μl TE, pH = 7 and submitted for sequencing. Samples were validated by TapeStation (Agilent) and qPCR before sequencing by Illumina^®^ MiSeq™. A ϕΧ174 viral genome library was routinely spiked in to the pooled sequencing samples because the low diversity of the target library would otherwise make it difficult to differentiate among clusters in the sequencing flowcell during the first few rounds of sequencing.

### Sequencing data analysis

Sequencing data analysis was performed using a custom code package written in MATLAB (MathWorks^®^), as described in the Results and Supplementary data. A comprehensive plain-language annotation of the code is provided in the Supplementary data.

Average per-base error frequencies (*f_error_*) based on the fraction of reads filtered (*R_filt_*) during a given step in data analysis were calculated by assuming a uniform average error rate across the length (in nucleotides, *L*) of the region being filtered such that *f_error_* = 1 − *f_correct_*; (*f_correct_*)*^L^* = (1 − *R_filt_*).

## RESULTS

### A self-priming hairpin construct

Each nonenzymatic RNA primer extension reaction involves two sequences of interest: the extended primer product sequence and the template sequence. We are interested in studying primer extension across randomised templates so we can better understand how the reaction accesses sequence space. If the reaction were error-free, then sequencing each product would automatically report on the corresponding complementary template. Similarly, if the template is defined, then sequencing each product is sufficient to interpret errors because deviations from complementarity to the defined template can be inferred as arising from mismatches (31). However, primer extension is not error-free and we seek to use randomised (not defined) templates. Therefore, we require an approach to sequencing that can supply both the primer extension product sequence and the corresponding template sequence.

One strategy to enable the extraction of product and template sequences is to physically connect the primer to the template with a hairpin loop so that the construct primes itself (Figure 1A) (36–38). Each sequencing read will then contain the template, an intervening defined hairpin, and any primer extension product sequence. We considered a variety of potential sequencing technologies and chose Illumina^®^ MiSeq™ because of its low error rate, forward and reverse read functionality (see below), and capacity to process multiple experimental samples simultaneously (multiplexing) through barcoding (indexing). Barcoding labels each sample with a unique identifier sequence so that multiple samples can be pooled and sequenced in one sequencing flowcell. The barcodes are later used to sort the data based on sample source. Preparing an RNA hairpin construct for deep-sequencing involves modifying it with additional sequences (Figure 1A-E), reverse transcription (Figure 1E-G), and barcoding through PCR (Figure 1G-I). The Illumina^®^ platform requires the addition of defined sequences (handles) to both 5’ and 3’ ends of the target. In our case the 3’ handle is also the binding site of the reverse transcription primer (RT Handle). Both handle sequences are used during barcoding PCR, and the PCR products include binding sites for the Illumina^®^ forward, reverse and barcode-reading sequencing primers.

**Figure 1.**
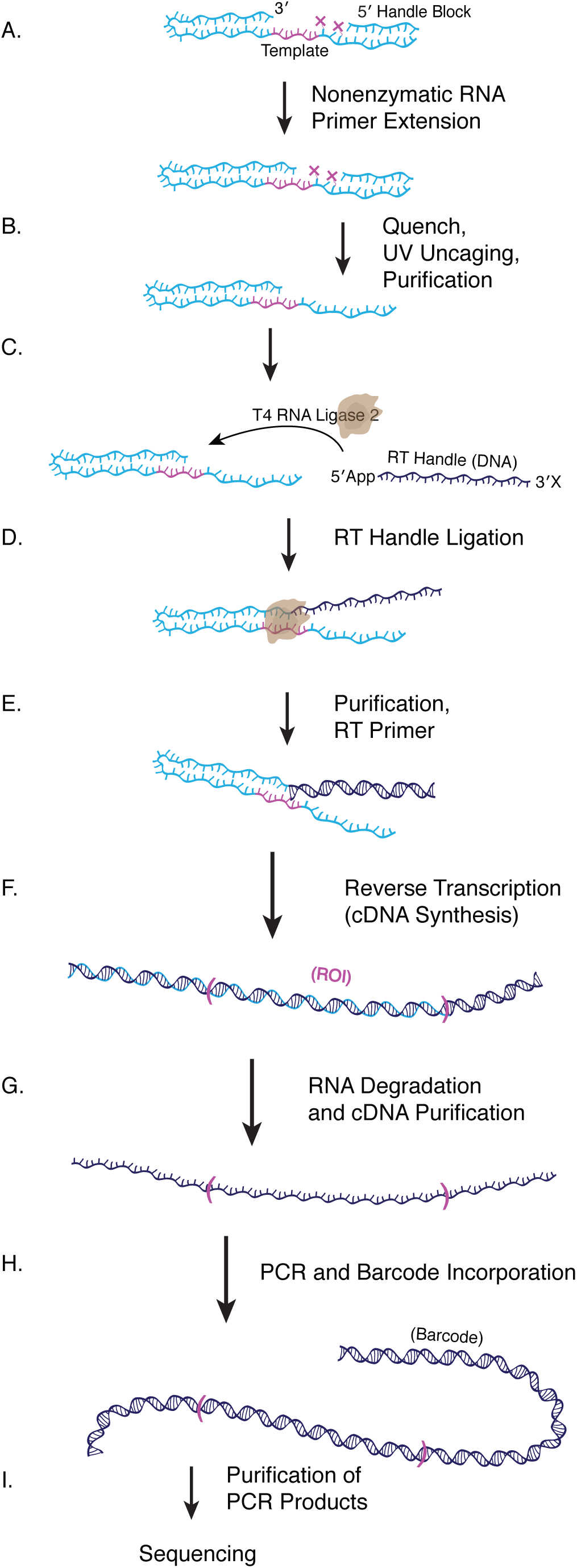
Protocol for Preparing RNA Hairpin Constructs for Sequencing. **A.** NERPE-Seq RNA hairpin constructs contain a hairpin loop that physically connects the template sequence to the primer so that both the product of nonenzymatic primer extension and the corresponding template are on one continuous RNA strand and can therefore be sequenced together. Two caged bases (magenta Xs) prevent primer extension from encroaching on the downstream 5’ Handle. The 5’ Handle Block is complementary to the 5’ Handle and prevents it from interfering with primer extension. **B.** The primer extension reaction is quenched with a desalting size-exclusion spin column, the caged bases are uncaged and the target RNA is further gel purified. **C-D.** The pre-adenylated DNA RT Handle (blocked on its 3’ end to prevent self-ligation) is ligated to the 3’ end of the RNA hairpin (the site of primer extension). **E.** The ligase is removed by a Proteinase K digestion, the target RNA-DNA is phenol-chloroform extracted, and the reverse transcription primer is annealed to the RT Handle. **F-G.** Reverse transcription generates the cDNA; the RNA is degraded, and the cDNA is isolated with a spin column. The Region of Interest (ROI) harbours the template, hairpin and any product sequences. **H.** PCR is used to barcode the DNA and add additional required flanking sequences. Each barcode identifies DNA from a specific experiment and enables the sequencing of samples drawn from multiple experimental conditions at the same time. **I.** The target PCR products are purified, and validated by automated electrophoresis and quantitative PCR prior to sequencing.

As a preliminary test of whether a hairpin construct could be effectively prepared for deep-sequencing, we adapted a standard RNA-Seq protocol in which the RT Handle is a 5’-adenylated (App), 3’-blocked DNA oligo ligated by an App ligase (a derivative of T4 RNA Ligase 2), and the 5’ Handle is ligated by T4 RNA Ligase 1 (39, 40). To test whether the ligases exhibit biases due to the hairpin secondary structure or varying extents of primer extension (41–43), we designed three mock constructs (Mock 1-3) with the desired self-priming structure, an eight-base “template” and varying extents of apparent primer extension (0, +3 and +7, respectively) (Figure S2). RT Handle ligation worked well on Mock 1 and Mock 2, but was less efficient on Mock 3, a mimic of substantial primer extension that is almost blunt, with only a single 5’ rA overhang (Figure S2A). Off-target products are also visible for Mock 3. The subsequent 5’ Handle ligation shows a similar pattern, with Mock 3 ligating least efficiently and showing additional off-target products (Figure S2B). Furthermore, the 5’ ligation was inefficient overall—in contrast to the RT Handle ligation— with most of the input material remaining unligated. The lower ligation efficiencies as a function of apparent primer extension length would be particularly problematic for interpreting data because substantially-extended products would be underrepresented. To compound these issues, the barcoding PCR of the ligation products yielded only primer dimers for Mock 3 (Figure S2B). We conclude that the standard RNA-Seq protocol is inadequate because it introduces side products and a pronounced bias against longer nonenzymatic primer extension products.

To begin addressing the problem of obtaining sequencing data that is an unbiased representation of primer extension, we noted that for our application the 5’ Handle sequence could be included as a component of the initial construct, thereby completely eliminating the need for a second ligation. However, the additional sequence downstream of the template introduces other challenges. First, if primer extension spans the template and then encroaches on the 5’ Handle sequence, the resultant product could interfere with sequencing and possibly complicate data analysis. Second, the additional single-stranded RNA of the 5’ Handle could interfere with primer extension by physically interacting with the template. Addressing the first challenge requires a feature that stops primer extension at a defined position without otherwise affecting the reaction, but does not inhibit reverse transcription or can be removed prior to reverse transcription. We chose 6-nitropiperonyloxymethyl (NPOM) amino caged deoxythymidine, an otherwise standard dT modified (“caged”) on the N3 of the base with the chemically stable NPOM moiety that can be removed (“uncaged”) by exposure to long-wavelength UV irradiation (44), outside the absorption spectrum of RNA (34). NPOM caging has the added benefits of inhibiting base pairing, especially if multiple NPOMs are used, and commercial availability as a phosphoramidite for solid-state oligo synthesis. (Inhibited base pairing is advantageous because it should limit interference of the 5’ Handle sequence with the templating region (though see below). It will also prevent short oligos used in some primer extension experiments (45) from binding to the 5’ Handle sequence and affecting primer extension; ideally, only the template sequence should affect the outcome of primer extension.) We prepared an RNA test oligo with a pair of NPOM-caged dT bases separated by an rC (Figure S3A) and exposed it to 385 nm UV light. PAGE analysis shows a gel shift after uncaging (Figure S3B), and LC-MS identified UV exposure conditions under which most of the oligos are uncaged (Figure S3C). We next tested whether the caged bases stop primer extension but allow reverse transcription when uncaged, and we found that the caged oligo induces an absolute stall on primer extension prior to the first caged dT, but allows full reverse transcription after uncaging (Figure S3D-F). Encouraged by the effectiveness of NPOM caging, we designed and synthesized a series of control hairpin constructs to facilitate protocol development (Figure S3G-J). The Control Template Extended construct (CTEx) mimics full-length primer extension on a defined template. The Control Template (CT) and Control Template B (CTB) constructs harbour a defined 3’-GCC-5’ template sequence, which is known to work well for primer extension (14). The 6N construct is the putative experimental substrate: it harbours six randomised template bases, representing all possible 6-mer templates. All constructs include a placeholding rA at the 5’ end of the template followed by two non-tandem caged bases.

#### Optimising RT handle ligation

Having defined a set of constructs we returned to the RT Handle ligation step, seeking to optimise it in the context of the encoded 5’ Handle sequence and NPOM caging. Noting that a base-paired 3’-end may be difficult to access for an enzyme, we tested a thermostable ligase. We reasoned that at higher temperatures the target 3’-end will breathe and thus possibly facilitate ligation, but the thermostable ligase is completely inactive for RT Handle ligation to 6N (Figure S4A). This may result from the particular hairpin structure of the construct and the inability of this particular enzyme to act on it. We also considered options for the adenylation and blocking of the RT Handle (Figure S4B-C). 5’ pre-adenylation of a handle increases the efficiency and specificity of ligation (5’ adenylate is an intermediate of the T4 RNA Ligase 2 enzymatic pathway (46)), and a blocked or absent 3’-OH prevents handle self-ligation. Ultimately, our optimised protocol employs a 3’ dideoxy, 5’ pre-adenylated RT Handle and a derivative of T4 RNA Ligase 2 (T4 RNA Ligase 2, truncated KQ (47)). Routine PAGE analysis during protocol development showed that this combination of ligase and handle works well across constructs. An ideal RT Handle ligation step would be equally efficient for all primer extension products while any hints of differential ligation, as previously observed for Mock 3 (Figure S2A), would bias the proportions of specific products in sequencing data. To optimise the ligation conditions we leveraged a particular case in which the reaction was found to be inefficient: RT Handle ligation to the +3 products of nonenzymatic trimer ligation to Control Template B (CTB) (Figure S4D-E). In this positive control nonenzymatic ligation reaction, a 2AI-activated 5’-CGG-3’ (2AI-CGG) trimer is incubated with the CTB construct, which has a complementary 3’-GCC-5’ template. We used this case to screen RT Handle ligation conditions and found that a higher incubation temperature (25°C instead of 16°C) combined with either DMSO or excess PEG_8000_ yielded complete RT Handle ligation to all reaction products to within the resolution of PAGE analysis. Higher temperature and DMSO are expected to destabilise the hairpin whereas PEG functions as a crowding agent, presumably favouring enzyme-substrate interactions. We chose the excess PEG_8000_ condition for inclusion in the final protocol, but note the potential usefulness of DMSO for ligation reactions involving RNA with a base-paired 3’ end.

### Preparing hairpin constructs for deep-sequencing

#### 2’-5’ linkages and reverse transcription

With a set of test constructs and optimal RT Handle ligation conditions in place, we next sought to characterise the reverse transcription (RT) step (Figure 1E-F). We considered three possible RT issues that might bias the final sequencing data: the uncaged bases, secondary structure inhibition and 2’-5’ linkages in the products of primer extension. RT had already been shown to work normally once the NPOM caging is removed (see Figure S3F). The uncaged residues are dT bases in an otherwise RNA template, and reverse transcribe to dA, as confirmed later by the sequencing data (see below). RNA-Seq experiments routinely process RNAs with secondary structures and a short hairpin stem should not impede reverse transcription. Hairpin-mediated stalling would register during PAGE analysis of reverse-transcribed products, but we do not find this to be an issue (see Figures 2 and S7). Previous research has shown that a single 2’-5’ linkage can be read by reverse transcriptase (48), but given our interest in mismatches, which exhibit a higher proportion of 2’-5’ linkages (23, 32), we tested whether multiple 2’-5’ linkages can also be read through by RT. We began with a test oligo harbouring a pair of 2’-5’ linkages separated by two bases (Figure S5A). Surprisingly, ProtoScript^®^ II reverse transcriptase is almost completely stalled by this substrate. However, the small amount of full-length DNA product (1.3%) suggests that although the enzyme is inefficient at reading through the 2’-5’ linkages, it is at least capable of doing so. This reaction was therefore used to screen conditions with the goal of maximising full-length reverse transcribed product (Figure S5B). Increasing the MgCl_2_ concentration from 3 mM to 10 mM combined with an overnight incubation at 42°C resulted in much better product yields, though stalling remained apparent (68% stalled and 32% full-length product) (Figure S5C). Importantly, these conditions do not degrade the RNA template (Figure S5D). We next prepared a series of RNA templates with increasingly separated pairs of 2’-5’ linkages (Figure S5E). As previously reported, a single 2’-5’ linkage does not significantly stall RT (90% full-length product) (48). Predictably, the most severe stall was measured for immediately adjacent 2’-5’ linkages (41% full-length product), whereas six intervening bases reduce the stalling effect (55% full-length product). A 2’-5’ linkage immediately adjacent to the primer consistently caused a stall regardless of the location of the subsequent 2’-5’ linkage. We conclude that primer extension products with multiple 2’-5’ linkages, especially at the first position after the RT primer or positioned tandemly, will be underrepresented in the sequencing data. Therefore, reported proportions of mismatches will represent a lower limit.

**Figure 2.**
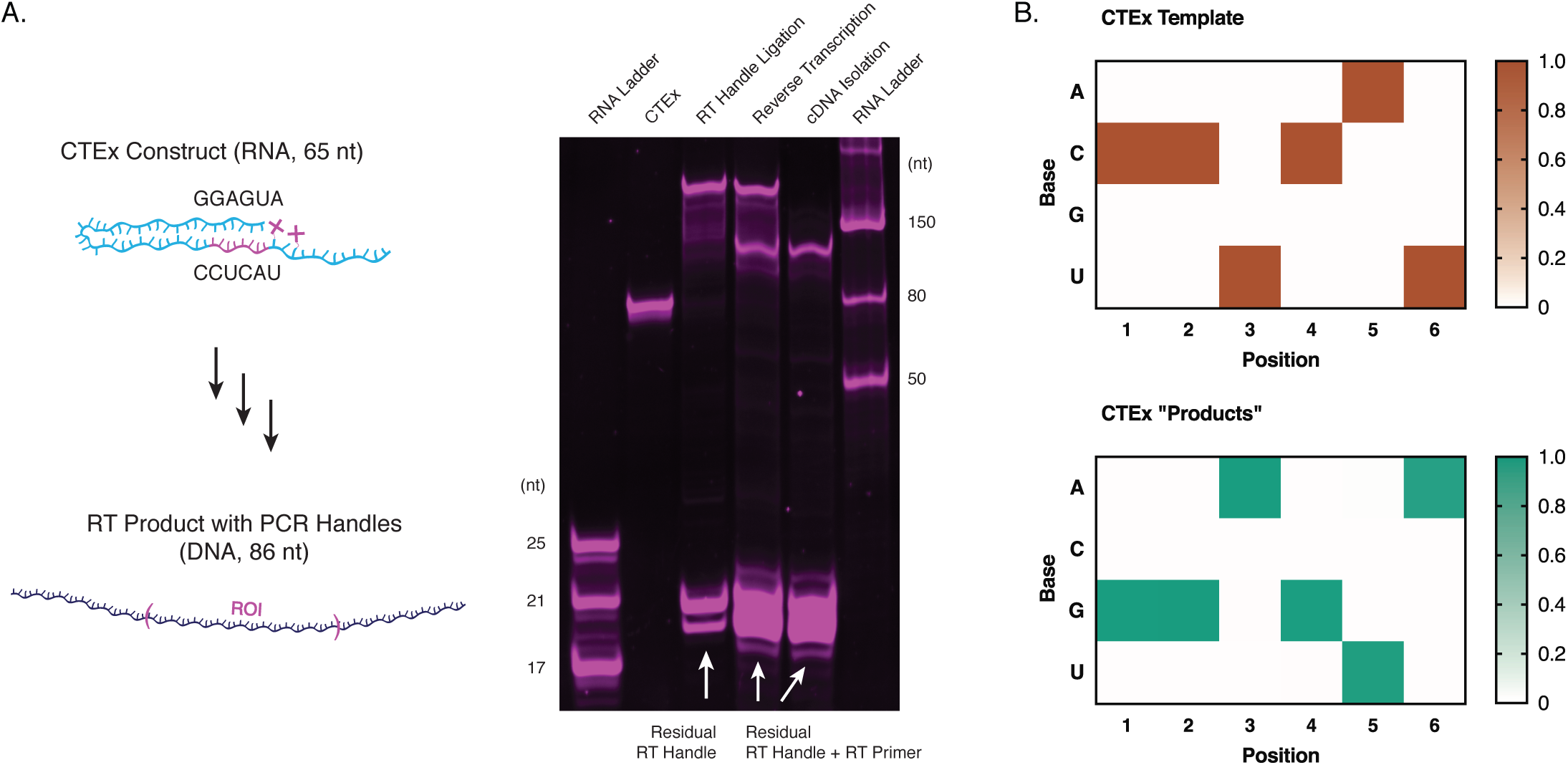
Sequencing a Construct that Mimics Full-length Nonenzymatic Primer Extension. **A.** A hairpin construct (CTEx) was designed to mimic an efficient nonenzymatic primer extension reaction. The RNA was converted into cDNA using the optimised protocol. The Region of Interest (ROI) includes the template and “product” sequences. After exposure to mock primer extension conditions and desalting, CTEx ran as a well-defined band at the expected position in PAGE. After uncaging and additional purification, the RT Handle was ligated to the 3’ end with very high efficiency. The product was purified and reverse transcribed, yielding a distinct DNA band, which was then isolated. **B.** The NERPE-Seq protocol accurately measures the CTEx template and “product” sequences without the template or product identities being provided during analysis. The heat maps show the frequency of each base at the indicated position.

#### Protocol trials and PCR optimisation

To evaluate the steps of the protocol established above—up to cDNA synthesis and isolation—by PAGE analysis, we used the Control Template Extended (CTEx) construct, which mimics efficient primer extension on a defined template (Figure 2A). CTEx was prepared under conditions equivalent to that used for primer extension experiments, incubated for 24 hours at 23°C and desalted. The CTEx RNA runs as a well-defined and undegraded band at the expected size after this treatment. It was then uncaged by UV irradiation, PAGE-purified, the RT Handle ligated and the product purified by Proteinase K treatment, phenol-chloroform extraction and spin column purification. The RT Handle ligation was extremely efficient. The ligated product was then reverse transcribed, yielding a distinct cDNA product band that runs at a lower mass than the template RNA. Although these two bands cannot be fairly compared directly because the signal intensity from the staining dye is not expected to be quantitative and the dye does not stain DNA and RNA equivalently, to a first approximation RT goes to completion. Finally, the RNA was degraded and the cDNA isolated by spin column purification, yielding a well-defined band of pure material. Notably, although CTEx mimics very efficient nonenzymatic primer extension, it exhibits none of the biases identified with the Mock 3 construct and the non-optimised protocol (compare Figure 2A with Figure S2A-B).

The final steps prior to sequencing submission are PCR barcoding and the isolation of target PCR products. In this protocol PCR does not serve to amplify the input material, which is not limiting, but rather to append additional flanking sequences and sample-specific barcodes. Therefore, the optimal PCR conditions maximise the input material (to limit the number of required cycles), minimise the number of cycles (to limit PCR artefacts) and minimise the concentration of PCR primers (to limit residual primers). Residual primers are especially problematic for deep-sequencing because they can lead to off-target amplification products and primer dimers. They can also directly contribute to barcode hopping, which is when a sequencing read is associated with the wrong barcode. One of the PCR primers carries the assigned barcode that uniquely identifies each sample and enables multiplexing. Residual primers that are introduced into the sequencing flowcell can randomly add their barcodes to DNA meant to exhibit a different barcode already assigned by PCR; this later results in the incorrect assignment of data during sequence analysis (barcode hopping, Illumina^®^ whitepaper 770-2017-004-D). A series of titrations identified the optimal PCR conditions as 2 ng/μl of input material, six cycles of amplification and 0.2 μM of each primer (Figure S6).

With the PCR optimised, we prepared a series of trials using the Control Template B (CTB) construct (Figure S7). As a negative control the CTB construct was exposed to primer extension conditions but without activated nucleotides so that no extension products were expected in the sequencing data (CTB Control, CTBC). As a positive control for canonical primer extension CTB was incubated with 20 mM each (40 mM total) of 2AIrG and 2AIrC (CTB 40mM activated nucleotides, CTB40). As a positive control for nonenzymatic ligation CTB was incubated with 500 μM 2AI-CGG trimer (CTB Ligation, CTBL). PAGE analysis was used to evaluate key steps of the protocol for each experiment, as for CTEx above, and all three show the expected banding patterns (Figure S7A). They also yielded well-defined PCR products as measured by agarose gel electrophoresis, as did CTEx (Figure S7B).

Samples submitted for sequencing are routinely quantified by automated electrophoresis, which is more sensitive than traditional agarose gel analysis. This enabled us to test a variety of target PCR product isolation strategies in pursuit of eliminating as much of the excess primer and any off-target PCR products as possible. We compared a PCR purification spin column, a magnetic bead-based purification and agarose gel purification (Figure S7C, 1-3). The spin column was largely ineffective at eliminating off-target bands, magnetic beads eliminated lower molecular weight off-target bands, and agarose gel purification eliminated higher molecular weight off-target bands. We therefore combined the magnetic bead and agarose gel purifications to yield very well-defined target material (Figure S7C, 4). Satisfied with the optimisation of this final step, we were prepared to submit samples for deep-sequencing and began formulating a toolkit to analyse the raw data.

### Data processing and analysis

The unique abiological nature of sequencing data from nonenzymatic RNA primer extension means that pre-existing software is not useful for answering the questions we are interested in, or even handling the raw data. For example, there is no reference genome, or any gene-based sequence structure to identify. We therefore developed a custom code package implemented in MATLAB (MathWorks^®^) to transform raw sequencing reads into a useful format (Pre-processing) and then make a series of key measurements (Characterisation) (Figure 3).

**Figure 3.**
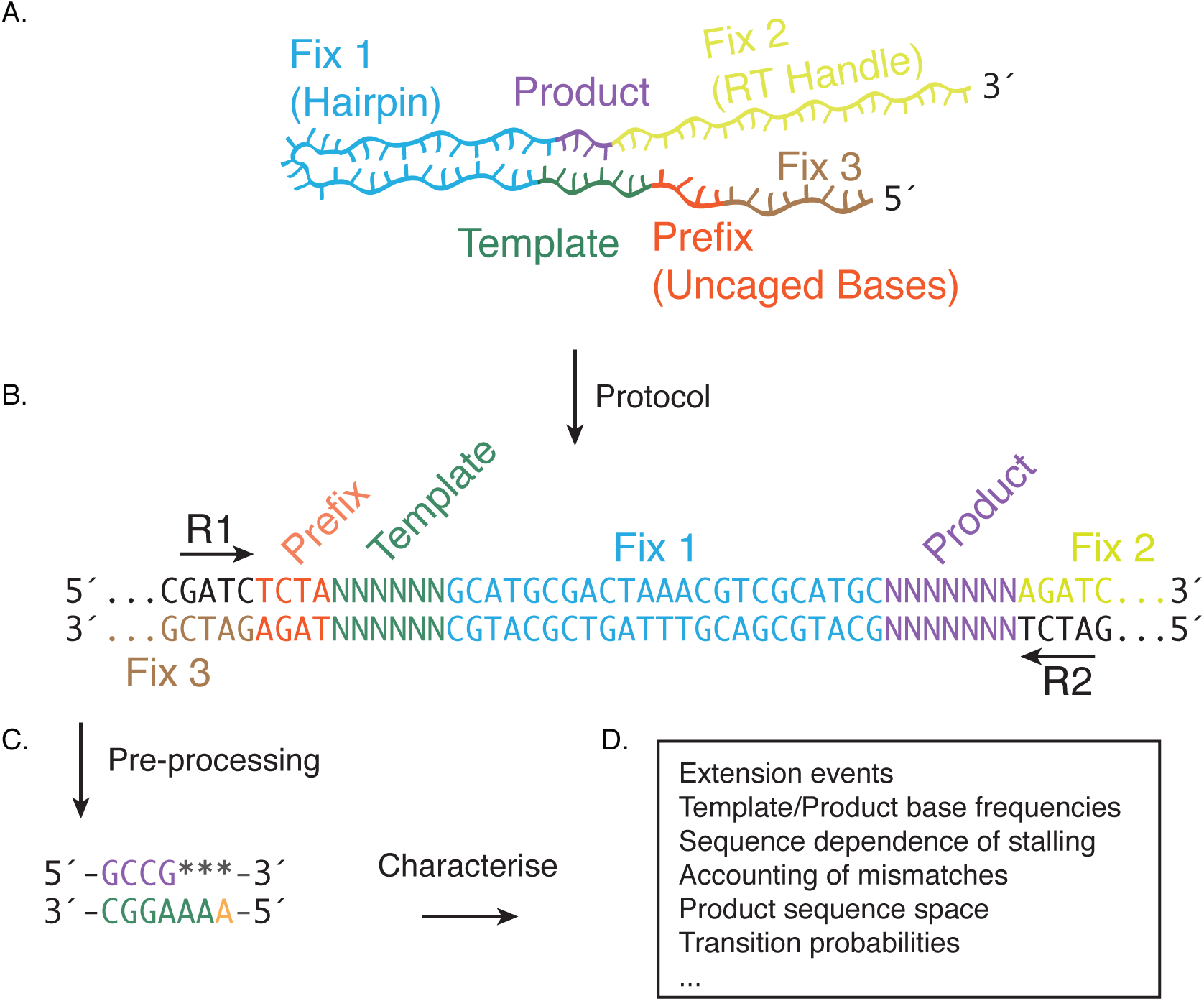
Data Analysis. **A.** Cartoon of a hairpin construct after RT Handle ligation. The labels and color coding indicate the various sequence regions used during data processing. The hairpin and handles are defined sequences (they are “fixed”), the Prefix is a four-base motif with the two caged bases, the Template is of defined length but the analysis does not specify a defined sequence (so we can analyse randomised templates), and the Product is of indeterminate length and sequence. **B.** The double-stranded DNA generated by barcoding PCR (the barcode is to the 3’ of Fix 2 and not shown). The final location of each region from the construct is labelled and color-coded. Paired-end sequencing provides both the forward (R1) and reverse (R2) sequences, which can be compared against each other for quality control. A series of checks identifies the fixed sequences, filters out low-quality reads, and extracts the Template and Product. **C.** The end result of Pre-processing is a set of Template-Product pairs. Unextended bases are indicated by an asterisk, and a placeholding A (part of the Prefix motif, immediately prior to the first caged base) is included as an internal marker. **D.** Template-Product pairs are queried in the Characterise stage to assay the sequence properties of templates and complementary products, and indicate the position and context of mismatches.

#### Pre-processing

This stage converts raw sequencing reads into a set of high quality template-product pairs, each representative of a template-primer complex from a single hairpin molecule (Figure 3A). We use paired-end sequencing so that each hairpin construct is sequenced twice: once in the “forward” direction (Read 1, R1), and once in the “reverse” direction to yield a reverse complementary sequence (Read 2, R2) (Figure 3B). R1 and R2 are compared to each other throughout Pre-processing to increase the quality of the output data. Raw sequencing files arrive as debarcoded sets in FASTQ format. (The MiSeq™ software automatically assigns each pair of forward and reverse reads to its associated barcode.) The reads in the files are broken up into blocks (typically 10,000 reads/block) to enable random access and parallel processing. Then, each read pair is subjected to a series of rigorous quality checks, and if it passes then the template and product (if any) sequences are extracted and stored.

The first quality check is to simply confirm that the headers (metadata and identifiers) match for R1, R2 and index (*i.e.*: barcode) read files. This ensures that only correctly paired reads are compared. The second check is based on estimated sequence accuracy. Estimated per-base Phred (49, 50) quality scores, *q*, provided by the sequencing run, are converted to per-base error probabilities, *p*, (*p* = 10^−*q*/10^) averaged across the read (*p̄* = *mean*(*p*)) and transformed back to an overall Phred quality score for the read, *q_p̄_* = −10*log*(*p̄*). A default threshold of *q_p̄_* = 26 was selected for the index (51), and *q_p̄_* = 30 for R1 and R2, which corresponds to a per-base error probability of 0.1% (1 in 1000).

The third quality check is correlative. We assume that a sequencing read with an error in one position is likely to have other errors. We therefore demand that sections of a read derived from defined (fixed) sequences in the hairpin construct (Fix 1-3 and Prefix) are correct and in the right location. Fix 1 corresponds to the hairpin sequence, Fix 2 to the RT Handle, Fix 3 to the 5’ Handle and Prefix to the “TCTA” motif derived from the two caged dT bases (Figure 3A). Forward reads begin with the Prefix and reverse reads begin with any products followed Fix 1 (Figure 3B). Crucially, identifying the fixed sequences enables us to identify the locations of the Template and Product. These can be of variable sequence (Template and Product) and variable length (Product), unlike the fixed sequences. We demand that R1 contain a single perfect match to Fix 2, which is only found in R1. We use this match to trim the end of R1 where Fix 2 starts. We next demand a perfect Fix 1 in both R1 and R2. The Template and Prefix are extracted based on their known lengths, and the Prefix is verified as matching its known sequence in both R1 and R2. We also demand that the Prefix location start at position 1 in R1, and that Fix 3 is present downstream of Prefix in R2. Then, we demand that the inferred product length make physical sense (cannot exceed the number of templating bases) and is equal in both R1 and R2. Finally, we demand that all of the trimmed R1 and (complement of) R2 agree, and generate the final Template-Product pair in RNA space (T → U) (Figure 3C). Each pair is oriented as in the source hairpin construct to intuitively reflect primer extension, with the primer extension product shown 5’-to-3’ on a 3’-to-5’ template, and the data are printed to a FASTA-like file.

If any step fails the read pair is discarded. For a typical multiplexed MiSeq™ run with eight barcoded samples, Pre-processing discards approximately half the initial data and yields approximately half a million read pairs per sample. For our purposes, this still supplies extremely high coverage (see below). Collectively, these stringent checks curtail barcode hopping, eliminate low quality reads, filter out low frequency artefacts that may be produced during PCR amplification or amplification on the flowcell, and provide high confidence in the accuracy of the template and product sequences.

#### Characterisation

This stage makes useful measurements on the set of high quality template-product pairs generated during Pre-processing. First, products of different lengths are quantified as position-dependent counts and frequencies. The resultant histogram is equivalent to the data accessible by PAGE analysis (see below). Next, we measure the position-dependent base distribution of the template. For a truly randomised template, each of the four bases would be equally represented at each position in a 0.25 proportion. However, it is difficult if not impossible to achieve such a perfect composition by solid-state oligonucleotide synthesis (see below and Figure S9A and B). We therefore calculate position-dependent template normalisation factors from the base distribution of the template. To eliminate template bias associated with a particular experimental condition (for example, an excess of 2’-5’ linkages), we include a comparison to template frequencies from a negative control experiment in which no primer extension is expected.

To facilitate the analysis of primer extension products, we group them into three non-overlapping classes: those with no extension events (Unextended Set, UnSet), those with extensions that are perfectly complementary to the template (Complementary Set, CompSet) and those with extensions that contain one or more mismatches (MisMatch Set, MMSet). We further characterise the position-dependent base composition of the CompSet products on an absolute, relative and normalised (to template or control template) basis.

To characterise the sequence-dependence of non-incorporation (*i.e.*: the termination of primer extension), we compute the position-dependent distribution of terminal product base counts and frequencies for the Complementary Set. To complement this measure and assess the effect of downstream templating bases on disfavouring extension, we also compute the position-dependent base distribution of all templating bases one base downstream of each terminal product. Unextended products are included because by definition their first templating position is one base downstream of a terminal (unextended) base. These counts are normalised to the template to account for template bias and expressed as per-position frequencies to allow comparison across positions.

As described above, primer extension proceeds via a bridged dinucleotide intermediate that binds to the template through Watson-Crick base pairing (Figure S1). We anticipate that adjacent pairs of templating bases in sequences with products will reflect the identity of the bridged dinucleotides that participated in primer extension. We therefore characterise the position-dependent pairs of adjacent bases that have corresponding correctly-paired products (that is, they belong to the Complementary Set). As this data cannot be reasonably normalised, we instead measure the position-dependent position-dependent pairs of adjacent bases across all templates in the sample for comparison. This measure is expected to directly correlate with the efficiency of primer extension as a function of bridged intermediate identity.

Identifying mismatches is a major goal of using deep-sequencing to study primer extension. The fidelity of primer extension (or, conversely, the error rate) is important for evaluating conditions under which emergent genetic information could be accurately propagated. In the MisMatch Set, we characterise the position-dependent counts of mismatches and the distribution of mismatches that are terminal. This is expected to reveal condition-dependent trends in overall primer extension fidelity and the impact of mismatches on extension termination. We also specifically look at position-dependent recovery, in which a mismatched base is followed by a correctly paired (complementary) base. Finally, we count the occurrence of each possible type of mismatch at each position (AA, AC, AG, CA, CC, CU, GA, GG, GU, UC, UG, UU) and compute their template-normalised frequencies.

To assay the sequence space accessed by primer extension we count all position-dependent sets of trimer products (*i.e.*: the identity of all three-base stretches in the products). We perform the same analysis on the template for comparison, and on just the Complementary Set. Any differences between the results for all product types and just the Complementary Set will show the extent to which mismatches contribute to the sequence space accessed by primer extension. Finally, we compute sequential base transition frequencies for the product and template. Here, we define sequential base transition frequencies as the probabilities of each subsequent base identity given the current base. Collectively, these metrics amount to a broad and deep accounting of nonenzymatic RNA primer extension sequence space and will be most powerful in the context of a randomised template.

### Deep sequencing results

Because the ultimate experimental goal of NERPE-Seq is to routinely analyse data from primer extension reactions on random template sequences, the software does not assume a specific template sequence or any particular primer extension product. We can therefore test the NERPE-Seq pipeline by submitting our control constructs and reactions, which *do* have specific templates and products, to see if NERPE-Seq yields the expected results. We first tested the Control Template Extended construct (CTEx), which mimics primer extension on a defined template, and found that NERPE-Seq accurately measures both the defined template and product sequences (Figure 2B). Encouraged by this result, we next considered a control experimental scenario that included a primer extension reaction.

#### The encoded 5’ handle can interfere with NERPE

The defined-template construct Control Template (CT, Figure S3H) was used to compare the results of primer extension as measured by PAGE analysis with the distribution of products as measured by NERPE-Seq. Disappointingly, the two datasets do not agree, with +2 products overrepresented and +3 products underrepresented in the NERPE-Seq analysis (Figure S8A-B). Prompted by orthogonal results with distinct constructs that show extremely good agreement between PAGE analysis and NERPE-Seq (manuscript in preparation), we designed a slightly different construct with a longer template region and two caged bases instead of one (Control Template B, CTB, Figure S3I). However, the PAGE and NERPE-Seq datasets in this case are in even greater disagreement (Figure S8C-D). A primer extension reaction with a separate primer, as used for PAGE-based experiments, and a self-primed hairpin reaction as used for NERPE-Seq should work equally well. However, we considered the possibility that the hairpin does not fold correctly. We screened a variety of hairpin folding conditions coupled to primer extension, but all reactions yielded the same characteristic pattern by PAGE analysis (Figure S8E), suggesting that the hairpin adopts the same optimal conformation across tested folding conditions. Finally, we considered that for the specific template sequence used in CT and CTB (3’-GCC-5’), the 5’ Handle may interfere with the template region despite the caged bases. We therefore included an oligo complementary to the 5’ Handle sequence (5’ Handle Block Test) to a primer extension reaction with CTB, and the expected product pattern was restored (Figure S8F). We conclude that portions of the 5’ Handle can transiently base pair with the template and inhibit primer extension, but if the ssRNA in the handle is occluded by a complementary strand to form duplex RNA, then primer extension proceeds normally and without interference (Figure S8G).

#### NERPE-Seq reproduces known results and can report key measurements

With the insight that blocking the 5’ Handle prevents it from interacting with the template and interfering with primer extension, we again compared the extent of primer extension using the primer-template that mimics Control Template B (CTB Mimic) (Figure 4A), as measured by PAGE analysis, with the extent of primer extension using Control Template B *with the 5’ Handle Block* (Figure 4B), as measured by NERPE-Seq. The reactions were incubated for 24 hours with 20 mM 2AIrG and 20 mM 2AIrC. The PAGE analysis shows predominantly +3 products, with some +2 and +4 (overextended) products, as expected (Figure 4C). The NERPE-Seq analysis shows the same distribution (Figure 4D), in excellent agreement with the PAGE results. We next tested whether nonenzymatic ligation could also be measured correctly by NERPE-Seq. The CTB Mimic primer-template and the CTB hairpin with the 5’ Handle Block were each incubated for 24 hours with 500 μM 2AI-CGG. Again, the product distribution as measured by PAGE and the product distribution as measured by NERPE-Seq are in excellent agreement (Figure 4E and 4F). These results demonstrate that NERPE-Seq can accurately and precisely make the same measurements of primer extension as PAGE analysis, the current standard in the field. NERPE-Seq can also plot product base distributions for each position (Figure 4G and 4H). The products are dominated by 5’-CGG-3’, as expected for a 3’-GCC-5’ template.

**Figure 4.**
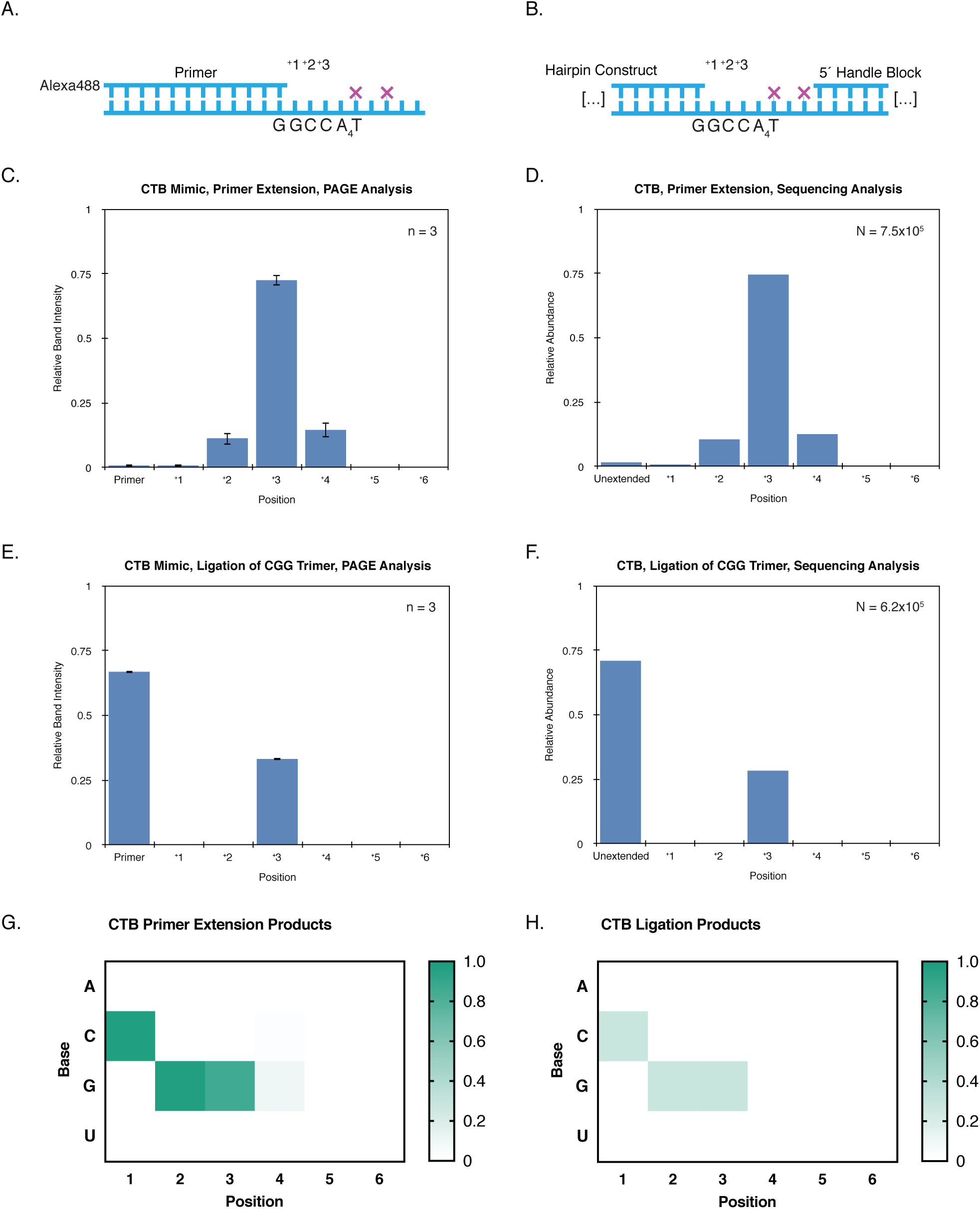
NERPE-Seq Can Accurately Measure Primer Extension on a Defined Template. **A.** CTB Mimic, used in PAGE analysis, is a construct that mimics the Control Template B (CTB) hairpin (**B**). **C.** The extent of primer extension with 20 mM each of 2AIrG and 2AIrC after 24 hours using CTB Mimic, as measured by PAGE analysis. **D.** The same reaction as in (C), but measured by NERPE-Seq on CTB. **E.** The extent of primer extension with 500 μM 2AI-CGG after 24 hours using CTB Mimic, as measured by PAGE analysis. **F.** The same reaction as in (E), but measured by NERPE-Seq on CTB. **G.** NERPE-Seq reveals the expected pattern of products from the polymerisation reaction (same experiment as in D) and (**H**) from the nonenzymatic ligation reaction (same experiment as in F). The heat maps show the frequency of each base at each position, including nulls. Note in (G) the mismatched overextension of G and C products across the templating A in position 4. In (H) the relative intensities at each position are equivalent because all products result from trimer ligation (*i.e.*: all ligation events contribute equivalently to the +1, +2 and +3 positions). Finally, the overall intensities in (G) are higher than in (H) because primer extension by polymerisation is in this case more efficient than ligation (as also reflected in the histograms: compare (D) to (F)).

In the case of Control Template B, errors in the defined template sequence can arise from oligo synthesis, whereas errors in the products cannot. The average per-base error in the template across four experiments with CTB is 0.12 ± 0.30%, which is within the range expected for oligo synthesis (correct per-base incorporation frequencies vary across oligo syntheses). The negative control experiment CTBC did not include any activated nucleotides; three additional experiments with CTB were incubated with 2AIrG and 2AIrC (see below). In these three experiments, incorrectly paired G or C products, which we expect (for example, a G product against the G in the first templating position) contribute 0.41 ± 0.46% per-base error. This is a measure of primer extension error in this specific experimental scenario. Products that are physically impossible (those that contain A or U, which were not added to the reaction) contribute 0.062 ± 0.13%: this is one estimate of our *experimental* per-base error because these products can only arise from sequencing errors, reverse transcription errors and barcode hopping, but not oligo synthesis or primer extension. The contribution of barcode hopping can be difficult to infer because it depends on the properties of the other samples in the sequencing run. To more directly measure barcode hopping, we performed a negative control experiment with the Control Template B hairpin (CTB Control, CTBC). In this experiment CTB was not exposed to activated nucleotides or oligonucleotides but was otherwise prepared and analysed exactly as the other samples. The vast majority of data from CTBC does not show any extension events, as expected, but 0.22% of the read pairs register as having at least one product base. This sequencing run contained seven other samples, all of which showed extensive primer extension. Assuming all the false positives resulted from barcode hopping, then each sample contributed 0.031% of the apparent extension products in this case. We take this as an estimate of the upper bound on barcode hopping. A more comprehensive accounting of error sources can be found in the Supplemental data.

Beyond the power to supply reliable data, we sought to develop an assay that can be applied across a variety of experimental conditions. To test whether we can monitor the consequences of distinct primer extension reactions, we performed a narrow titration of activated nucleotide concentrations using the defined-template Control Template B, from 10 mM each of 2AIrG and 2AIrC (CTB20), through 20 mM each (CTB40, described above), to 30 mM each (CTB60). We expect a lower proportion of +3 products for CTB20 than CTB40, which is what we observe: 70% for CTB20 versus 75% for CTB40. Interestingly, the proportion of +3 products for CTB60 does not follow this trend, with 67% +3, but also slightly more +2 and +4 than observed in the other two experiments. This suggests that increasing the activated nucleotide concentration is not necessarily efficient at driving primer extension to the ideal products. Future work will explore potential explanations for such phenomena. We also expect overextension to the +4 position, which harbours a templating rA, to rise with concentration. We observe that each increase in activated nucleotide concentration also increases overextension, with +4 extensions accounting for 7% of CTB20 products, 13% of CTB40 products and 16% of CTB60 products. Taken together, these results indicate that NERPE-Seq analysis is sensitive to changes in experimental conditions.

An essential application of NERPE-Seq is to characterise the occurrences of mismatches during nonenzymatic template copying. Mismatches were measured in the cases described above, but those experiments were not designed to specifically yield mismatches. We therefore performed an experiment in which Control Template B was incubated with 20 mM 2AIrU for 24 hours (CTB mismatch, CTBmm). All products should be U bases incorrectly paired across the 3’-GCC-5’ template, and this is what we observe (Figure 5A). As expected for a necessarily incorrect primer extension, the overall efficiency was much lower than in the experiments above, with only 6.1% of read pairs registering extension events. We can also visualise all position-dependent mismatches (Figure 5B). The G-U mismatch dominates position 1 and the C-U mismatch dominates position 2, both as expected. The additional apparent mismatches in downstream positions presumably result from sequencing errors. Notably, the vast majority of the data in this case (>98%) falls in the position 1 bin, with just under 1% in the position 2 bin. This suggests that mismatch data begins to encounter error noise when a given position harbours less than 1% of all mismatches, a useful metric for future analyses.

**Figure 5.**
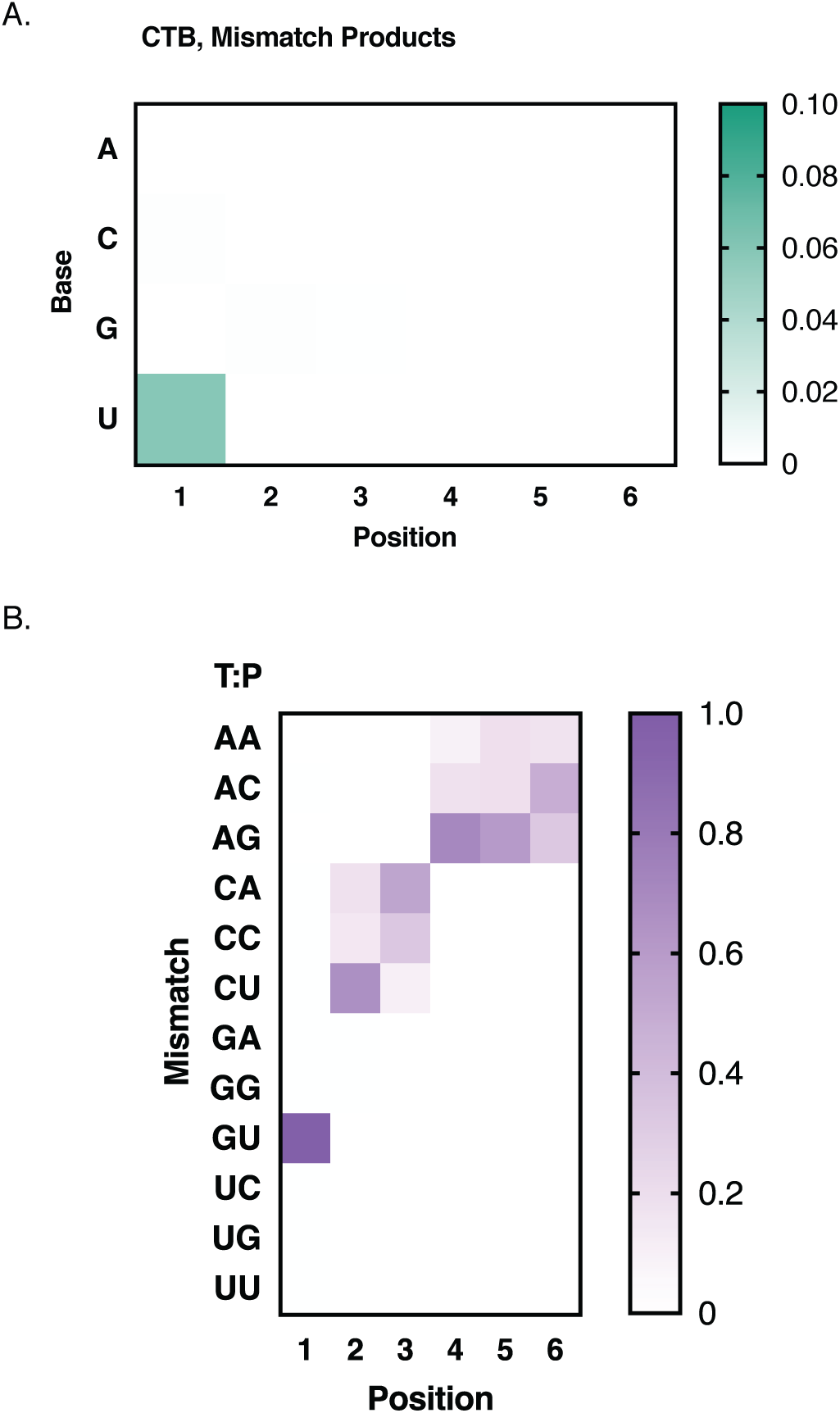
NERPE-Seq Can Measure Mismatch Frequencies. **A.** NERPE-Seq reveals the expected pattern of products from a primer extension reaction designed to intentionally generate mismatches. The Control Template B hairpin, which has a GCC template, was incubated with 20 mM 2AIrU for 24 hours. The heat map shows the proportion of each base at each position, including nulls. Note the intensity scale bar maximum value is set to 0.1 because there are relatively few products. **B.** The position-dependent distribution of all possible mismatches. The heat map shows the proportion of each mismatch at each position relative to all mismatches in that position. As expected, the Template:Product (T:P) mismatch of G:U in the first position dominates the data, and C:U is also evident in the second position, as expected. The data become noisy even by position 2 because there are very few extension events beyond +1 (>98% of the data is in position 1).

#### The 6N template and oligonucleotide synthesiser errors

Our data analysis assumes that a synthetic template designed to be random will not actually be truly random, and accounts for this deviation by calculating normalisation factors to correctly adjust product proportions (see above). The phosphoramidites used in solid-state oligo synthesis are known to exhibit distinct coupling efficiencies (52), but even if these are accounted for by mixing appropriately scaled molar quantities of each base, we cannot assume that the product will always be exactly random. This is important to understand for future experiments that will rely on constructs such as 6N, a hairpin construct with a 6-nucleotide random template prepared with an equimolar mixture of all four phosphoramidites. NERPE-Seq automatically measures the template base distributions, so we performed a negative control experiment in which 6N (6N Control, 6NC) was not exposed to activated nucleotides or oligonucleotides but was otherwise prepared and analysed exactly as the other samples. The template is not fully random (Figure S9A), as G is overrepresented and C is underrepresented. Much more surprising is that the base distribution changes as a function of position. To better understand this observation, we measured the position-dependent sequential base transition frequencies (the probability of each possible subsequent base given the current base, Figure S9B) and found a significant amount of structure. In particular, sequential transitions from C-to-G (that is, 3’-CG-sequences in the template) are favoured, and there are other more subtle, though consistent, effects. This suggests that truly random distributions by solid-state oligo synthesis are inherently impossible because pairwise base identities influence coupling efficiencies. These results highlight the necessity of the normalisation approach we implemented in our data analysis.

The data analysis filters out sequencing reads with an average quality score below 30, a threshold that corresponds to a per-base error rate of 0.1%. The analysis also checks reads for perfect matches to the fixed sequences from the hairpin construct. These checks typically flag about half of the data for disposal, which is significantly more than would be expected if only sequencing errors were involved. For example, 23.4% of read pairs that had already passed with the requisite average quality score were filtered out of the 6N data because they did not harbour a correct Fix 1 in R1. This corresponds to a 1.15% per-base error, much higher than expected after initial quality filtering. We suggest that this additional error is primarily due to imperfect oligo synthesis. This interpretation implies that the errors are genuine incorrect bases present in the sequence rather than mistakes made by the sequencer, and should therefore occur in both R1 and R2. (If they were sequencing errors, then the error frequency in R1 and R2 would be uncorrelated or only weakly correlated.) Recall that the data analysis filters sequencing reads in pairs: if an R1 read is filtered out, then the corresponding R2 read is also automatically discarded. Therefore, if an error occurs in both R1 and R2, then the R1 filter pass should eliminate the corresponding R2 reads. When R2 is subsequently filtered for correct Fix 1 sequences, only an additional 0.021% of read pairs are discarded, *corresponding to an average per-base error of only 0.00089%* (one estimate of sequencing error). This strongly suggests that Fix 1 is sequenced correctly but contains errors from the synthesiser: read pairs with deviations from the known Fix 1 sequence are discarded during the R1 pass, and the R2 pass only results in a small additional loss due to sequencing errors. (Barcode hopping is not relevant in this case because all constructs have the same Fix 1.)

To see whether synthesiser errors show any interpretable patterns we plotted the base distributions of Fix 1 errors from the filtered R1 reads (Figure S9C). Each contributing read contains at least one base that is incorrect compared with the defined Fix 1 sequence. The errors appear somewhat random overall, but in more than half of positions the highest frequency incorrect base is the same as the *preceding* correct base in the 3’-to-5’ direction, which is the direction in which the oligonucleotide was synthesised. The simplest explanation is contamination during a given synthesis cycle from the previously added phosphoramidite, which would result in a substitution mutation. Alternatively, one phosphoramidite base could be added twice in a single synthesis cycle, resulting in an insertion mutation. Both types of errors are filtered out by Pre-processing. We conclude that the large proportion of filtered read pairs results primarily from errors during oligo synthesis and that a discernible fraction of those errors can be traced to line contamination or double addition during synthesis cycles.

## DISCUSSION

Experimentally capturing the properties of an emergent life-like replicating system will require accommodating the complexity of reaction networks, environmental and chemical cycling and heterogeneity (25,28,53–56). A key facet of complexity in an RNA world scenario is the sequence space accessed by nonenzymatic copying. It is this sequence space that will determine how, or even whether, primitive RNA phenotypes can be explored. Although the chemistry of nonenzymatic polymerisation and ligation is now well-understood, much less is known about how it will influence copied sequence properties generally, especially in the context of random templates and oligomers and (at least) all four canonical ribonucleotides. Furthermore, the fidelity of primer extension will dictate whether favourable sequences can be maintained across cycles of copying. Mismatch identities and prevalence, and their sequence-dependence and reaction-condition-dependence are therefore essential to understanding how evolutionary selection may have emerged in the RNA world. Current methods are incapable of measuring the extent of sequence space accessed by nonenzymatic primer extension and comprehensively measuring mismatches under different conditions.

Nonenzymatic RNA primer extension experiments are traditionally assayed by HPLC or electrophoresis, especially denaturing PAGE. Both techniques separate products based on size, and interpreting the results requires knowledge of the template and potential product sequences. Mass spectroscopy can precisely identify multiple products if the number of possible outcomes is not so high that the spectrogram becomes too densely populated to interpret (32,57,58). Here too the template must be defined for the results to be unambiguous. HPLC, PAGE and mass spectroscopy work on non-canonical RNA (59–62), unlike sequencing, which relies on biological enzymes. Deep-sequencing has previously been used to characterise the products of nonenzymatic primer extension on a small set of defined templates (31). That study isolated the product-containing strands and inferred mismatch identities based on the known template sequences. To-date, there has been no generalisable method to directly measure individual template-product pairs across sequence space.

The NERPE-Seq analysis automatically generates comprehensive data about the templates, products and mismatches in a primer extension experiment, and multiplexing through barcoding PCR enables us to easily sequence samples from multiple experiments at once. We also gain the benefit of high throughput because deep-sequencing generates millions of sequencing reads. Quality filtering jettisons about half of the data, primarily because of errors from oligo synthesis. However, NERPE-Seq experiments are not currently data-limited. In the case of an RNA hairpin construct with six randomised template bases there are 4^6^ = 4,096 possible template sequences, so the approximately half million read pairs typically retained per sample after filtering affords over 100X coverage. In addition to the applicability of NERPE-Seq to any experimental condition compatible with RNA, the constructs are made by solid-state oligo synthesis and can therefore harbour any template sequence chosen by the experimenter, including random sequences of any length (within the limits of synthesis capabilities). This versatility enables the study of any standard RNA primer extension experiment on any template, and the use of substrates in any combination so long as they are compatible with reverse transcription (63). We anticipate that NERPE-Seq could easily be modified for DNA-based nonenzymatic reactions (64), as well as biological or putative ribozyme-catalysed primer extension (65).

As we begin to probe higher-order primer extension experiments, the set of reactants will only multiply. Experiments with all four nucleotides are rare, partly because the reaction is inefficient and partly because the results are challenging to interpret without sequencing data (56, 59). NERPE-Seq may help us address the relationship between inefficient primer extension with all four bases and the role of the bridged intermediate. A unique feature of the bridged intermediate pathway is that sequential pairs of templating bases are relevant to each additional added base. This means that the +2 templating position, and whatever substrates occupy it, or attempt to occupy it, can affect what happens at the +1 product position. NERPE-Seq will directly report on these features. Of similar interest is the role, if any, of the ratios of unactivated nucleotides, activated nucleotides and bridged dinucleotides on extension efficiency and fidelity. Finally, NERPE-Seq will be extremely powerful in assaying even more heterogeneous systems containing short oligonucleotides of different lengths (45,66,67) and various distributions of activated and inactivated material (28, 57). Such complex mixtures of reactants are more realistic than pure monoribonucleotides. Furthermore, oligos have been demonstrated to facilitate primer extension in defined sequence contexts, but it remains unknown how a randomised mixture, such as we would expect to find among a propagating pool of self-replicating RNAs, would change the efficiency and fidelity of primer extension. NERPE-Seq will enable the routine analysis of such experiments in outstanding detail.

## DATA AVAILABILITY

The NERPE-Seq analysis code is available in the GitHub repository: https://github.com/CarrCE/NERPE-Seq

Sequencing reads and NERPE-Seq analysis output data will be made available at OSF.io on formal publication.

## ACKNOWLEDGEMENTS

We thank the staff of the MGH Department of Molecular Biology Next-generation Sequencing Core for sample validation, MiSeq™ runs and communicating technical knowledge. We are grateful to Saurja DasGupta, Li Li, Stephanie Zhang, Lijun Zhou and Aleksandar Radaković for comments on the manuscript, and members of the Szostak laboratory for feedback and sharing experimental expertise.

## REFERENCES

1. Gilbert, W. (1986) Origin of Life - The RNA World. Nature, 319, 618–618.

2. Orgel, L.E. (1989) Was RNA the First Genetic Polymer? Evolutionary Tinkering in Gene Expression, 169, 215–224.

3. Joyce, G.F. (2002) The antiquity of RNA-based evolution. Nature, 418, 214–221.

4. Robertson, M.P. and Joyce, G.F. (2012) The Origins of the RNA World. Cold Spring Harbor Perspect. Biol., 4.

5. Krishnamurthy, R. (2015) On the Emergence of RNA. Israel Journal of Chemistry, 55, 837–850.

6. Szostak, J.W. (2017) The Narrow Road to the Deep Past: In Search of the Chemistry of the Origin of Life. Angew. Chem.-Int. Edit., 56, 11037–11043.

7. Joyce, G.F. and Szostak, J.W. (2018) Protocells and RNA Self-Replication. Cold Spring Harbor Perspect. Biol., 10.

8. Weimann, B.J., Lohrmann, R., Orgel, L.E., Schneide. H and Sulston, J.E. (1968) Template-directed Synthesis With Adenosine-5’-Phosphorimidazolide. Science, 161, 387–&.

9. Sulston, J., Lohrmann, R., Orgel, L.E. and Miles, H.T. (1968) Nonenzymatic Synthesis of Oligoadenylates on a Polyuridylic Acid Template. Proc. Natl. Acad. Sci. U. S. A., 59, 726–&.

10. Lohrmann, R., Bridson, P.K., Bridson, P.K. and Orgel, L.E. (1980) Efficient Metal Ion Catalyzed Template-directed Oligonucleotide Synthesis. Science, 208, 1464–1465.

11. Rohatgi, R., Bartel, D.P. and Szostak, J.W. (1996) Kinetic and mechanistic analysis of nonenzymatic, template-directed oligoribonucleotide ligation. J. Am. Chem. Soc., 118, 3332–3339.

12. Kervio, E., Sosson, M. and Richert, C. (2016) The effect of leaving groups on binding and reactivity in enzyme-free copying of DNA and RNA. Nucleic Acids Res., 44, 5504–5514.

13. Fahrenbach, A.C., Giurgiu, C., Tam, C.P., Li, L., Hongo, Y., Aono, M. and Szostak, J.W. (2017) Common and Potentially Prebiotic Origin for Precursors of Nucleotide Synthesis and Activation. J. Am. Chem. Soc., 139, 8780–8783.

14. Li, L., Prywes, N., Tam, C.P., O’Flaherty, D.K., Lelyveld, V.S., Izgu, E.C., Pal, A. and Szostak, J.W. (2017) Enhanced Nonenzymatic RNA Copying with 2-Aminoimidazole Activated Nucleotides. J. Am. Chem. Soc., 139, 1810–1813.

15. Walton, T. and Szostak, J.W. (2016) A Highly Reactive Imidazolium-Bridged Dinucleotide Intermediate in Nonenzymatic RNA Primer Extension. J. Am. Chem. Soc., 138, 11996–12002.

16. Walton, T. and Szostak, J.W. (2017) A Kinetic Model of Nonenzymatic RNA Polymerization by Cytidine-5’-phosphoro-2-aminoimidazolide. Biochemistry, 56, 5739–5747.

17. Zhang, W., Tam, C.P., Walton, T., Fahrenbach, A.C., Birrane, G. and Szostak, J.W. (2017) Insight into the mechanism of nonenzymatic RNA primer extension from the structure of an RNA-GpppG complex. Proc. Natl. Acad. Sci. U. S. A., 114, 7659–7664.

18. Zhang, W., Walton, T., Li, L. and Szostak, J.W. (2018) Crystallographic observation of nonenzymatic RNA primer extension. eLife, 7, 15.

19. Walton, T., Zhang, W., Li, L., Tam, C.P. and Szostak, J.W. (2019) The Mechanism of Nonenzymatic Template Copying with Imidazole-Activated Nucleotides. Angew. Chem.-Int. Edit., 58, 10812–10819.

20. Lohrmann, R. and Orgel, L.E. (1978) Preferential Formation of (2’-5’)-linked Internucleotide Bonds in Non-enzymatic Synthesis. Tetrahedron, 34, 853–855.

21. Rohatgi, R., Bartel, D.P. and Szostak, J.W. (1996) Nonenzymatic, template-directed ligation of oligoribonucleotides is highly regioselective for the formation of 3’-5’ phosphodiester bonds. J. Am. Chem. Soc., 118, 3340–3344.

22. Sheng, J., Li, L., Engelhart, A.E., Gan, J.H., Wang, J.W. and Szostak, J.W. (2014) Structural insights into the effects of 2’-5’ linkages on the RNA duplex. Proc. Natl. Acad. Sci. U. S. A., 111, 3050–3055.

23. Giurgiu, C., Li, L., O’Flaherty, D.K., Tam, C.P. and Szostak, J.W. (2017) A Mechanistic Explanation for the Regioselectivity of Nonenzymatic RNA Primer Extension. J. Am. Chem. Soc., 139, 16741–16747.

24. Powner, M.W., Gerland, B. and Sutherland, J.D. (2009) Synthesis of activated pyrimidine ribonucleotides in prebiotically plausible conditions. Nature, 459, 239–242.

25. Patel, B.H., Percivalle, C., Ritson, D.J., Duffy, C.D. and Sutherland, J.D. (2015) Common origins of RNA, protein and lipid precursors in a cyanosulfidic protometabolism. Nat. Chem., 7, 301–307.

26. Sutherland, J.D. (2016) The Origin of Life-Out of the Blue. Angew. Chem.-Int. Edit., 55, 104–121.

27. Xu, J.F., Tsanakopoulou, M., Magnani, C.J., Szabla, R., Sponer, J.E., Sponer, J., Gora, R.W. and Sutherland, J.D. (2017) A prebiotically plausible synthesis of pyrimidine beta-ribonucleosides and their phosphate derivatives involving photoanomerization. Nat. Chem., 9, 303–309.

28. Mariani, A., Russell, D.A., Javelle, T. and Sutherland, J.D. (2018) A Light-Releasable Potentially Prebiotic Nucleotide Activating Agent. J. Am. Chem. Soc., 140, 8657–8661.

29. Gibard, C., Bhowmik, S., Karki, M., Kim, E.K. and Krishnamurthy, R. (2018) Phosphorylation, oligomerization and self-assembly in water under potential prebiotic conditions. Nat. Chem., 10, 212–217.

30. Rajamani, S., Ichida, J.K., Antal, T., Treco, D.A., Leu, K., Nowak, M.A., Szostak, J.W. and Chen, I.A. (2010) Effect of Stalling after Mismatches on the Error Catastrophe in Nonenzymatic Nucleic Acid Replication. J. Am. Chem. Soc., 132, 5880–5885.

31. Heuberger, B.D., Pal, A., Del Frate, F., Topkar, V.V. and Szostak, J.W. (2015) Replacing Uridine with 2-Thiouridine Enhances the Rate and Fidelity of Nonenzymatic RNA Primer Extension. J. Am. Chem. Soc., 137, 2769–2775.

32. Leu, K., Kervio, E., Obermayer, B., Turk-MacLeod, R.M., Yuan, C., Luevano, J.M., Chen, E., Gerland, U., Richert, C. and Chen, I.A. (2013) Cascade of Reduced Speed and Accuracy after Errors in Enzyme-Free Copying of Nucleic Acid Sequences. J. Am. Chem. Soc., 135, 354–366.

33. Prywes, N., Michaels, Y.S., Pal, A., Oh, S.S. and Szostak, J.W. (2016) Thiolated uridine substrates and templates improve the rate and fidelity of ribozyme-catalyzed RNA copying. Chem. Commun., 52, 6529–6532.

34. Cavaluzzi, M.J. and Borer, P.N. (2004) Revised UV extinction coefficients for nucleoside-5’-monophosphates and unpaired DNA and RNA. Nucleic Acids Res., 32.

35. Schindelin, J., Arganda-Carreras, I., Frise, E., Kaynig, V., Longair, M., Pietzsch, T., Preibisch, S., Rueden, C., Saalfeld, S., Schmid, B. et al. (2012) Fiji: an open-source platform for biological-image analysis. Nature Methods, 9, 676–682.

36. Wu, T.F. and Orgel, L.E. (1992) Nonenzymatic Template-directed Synthesis on Oligodeoxycytidylate Sequences in Hairpin Oligonucleotides. J. Am. Chem. Soc., 114, 317–322.

37. Wu, T.F. and Orgel, L.E. (1992) Nonenzymatic Template-directed Synthesis on Oligodeoxycytidylate Sequences in Hairpin Oligonucleotides 2. Templates Containing Cytidine and Guanosine Residues. J. Am. Chem. Soc., 114, 5496–5501.

38. Wu, T. and Orgel, L.E. (1992) Nonenzymatic Template-directed Synthesis on Oligodeoxycytidylate Sequences in Hairpin Oligonucleotides 3. Incorporation of Adenosine and Uridine Residues. J. Am. Chem. Soc., 114, 7963–7969.

39. Ozsolak, F. and Milos, P.M. (2011) RNA sequencing: advances, challenges and opportunities. Nature Reviews Genetics, 12, 87–98.

40. Zhang, Z.J., Lee, J.E., Riemondy, K., Anderson, E.M. and Yi, R. (2013) High-efficiency RNA cloning enables accurate quantification of miRNA expression by deep sequencing. Genome Biology, 14.

41. Hafner, M., Renwick, N., Brown, M., Mihailovic, A., Holoch, D., Lin, C., Pena, J.T.G., Nusbaum, J.D., Morozov, P., Ludwig, J. et al. (2011) RNA-ligase-dependent biases in miRNA representation in deep-sequenced small RNA cDNA libraries. Rna-a Publication of the Rna Society, 17, 1697–1712.

42. Zhuang, F.L., Fuchs, R.T., Sun, Z.Y., Zheng, Y. and Robb, G.B. (2012) Structural bias in T4 RNA ligase-mediated 3’-adapter ligation. Nucleic Acids Res., 40.

43. Fuchs, R.T., Sun, Z.Y., Zhuang, F.L. and Robb, G.B. (2015) Bias in Ligation-Based Small RNA Sequencing Library Construction Is Determined by Adaptor and RNA Structure. Plos One, 10.

44. Lusic, H. and Deiters, A. (2006) A new photocaging group for aromatic N-heterocycles. Synthesis-Stuttgart, 2147–2150.

45. Prywes, N., Blain, J.C., Del Frate, F. and Szostak, J.W. (2016) Nonenzymatic copying of RNA templates containing all four letters is catalyzed by activated oligonucleotides. eLife, 5.

46. Nandakumar, J., Shuman, S. and Lima, C.D. (2006) RNA ligase structures reveal the basis for RNA specificity and conformational changes that drive ligation forward. Cell, 127, 71–84.

47. Viollet, S., Fuchs, R.T., Munafo, D.B., Zhuang, F. and Robb, G.B. (2011) T4 RNA Ligase 2 truncated active site mutants: improved tools for RNA analysis. BMC Biotechnology, 11, 72.

48. Lorsch, J.R., Bartel, D.P. and Szostak, J.W. (1995) Reverse-transcriptase Reads Through a 2’-5’-linkage and a 2’-thiophosphate in a Template. Nucleic Acids Res., 23, 2811–2814.

49. Ewing, B. and Green, P. (1998), Genome Res, Vol. 8, pp. 186–194.

50. Ewing, B., Hillier, L., Wendl, M.C. and Green, P. (1998), Genome Res, Vol. 8, pp. 175–185.

51. Wright, E.S. and Vetsigian, K.H. (2016) Quality filtering of Illumina index reads mitigates sample cross-talk. Bmc Genomics, 17.

52. Andrew Ellington and Jack D., Pollard, J. (2000) Introduction to the Synthesis and Purification of Oligonucleotides. Current Protocols in Nucleic Acid Chemistry, A.3C.1–A.3C.22.

53. Szostak, J.W. (2011) An optimal degree of physical and chemical heterogeneity for the origin of life? Philosophical Transactions of the Royal Society B-Biological Sciences, 366, 2894–2901.

54. Budin, I., Prwyes, N., Zhang, N. and Szostak, J.W. (2014) Chain-Length Heterogeneity Allows for the Assembly of Fatty Acid Vesicles in Dilute Solutions. Biophys. J., 107, 1582–1590.

55. Larsen, A.T., Fahrenbach, A.C., Sheng, J., Pian, J.L. and Szostak, J.W. (2015) Thermodynamic insights into 2-thiouridine-enhanced RNA hybridization. Nucleic Acids Res., 43, 7675–7687.

56. Mariani, A. and Sutherland, J.D. (2017) Non-Enzymatic RNA Backbone Proofreading through Energy-Dissipative Recycling. Angew. Chem.-Int. Edit., 56, 6563–6566.

57. Deck, C., Jauker, M. and Richert, C. (2011) Efficient enzyme-free copying of all four nucleobases templated by immobilized RNA. Nat. Chem., 3, 603–608.

58. Sosson, M. and Richert, C. (2018) Enzyme-free genetic copying of DNA and RNA sequences. Beilstein Journal of Organic Chemistry, 14, 603–617.

59. Blain, J.C., Ricardo, A. and Szostak, J.W. (2014) Synthesis and Nonenzymatic Template-Directed Polymerization of 2’-Amino-2’-deoxythreose Nucleotides. J. Am. Chem. Soc., 136, 2033–2039.

60. O’Flaherty, D.K., Zhou, L.J. and Szostak, J.W. (2019) Nonenzymatic Template-Directed Synthesis of Mixed-Sequence 3’-NP-DNA up to 25 Nucleotides Long Inside Model Protocells. J. Am. Chem. Soc., 141, 10481–10488.

61. Kim, S.C., O’Flaherty, D.K., Zhou, L.J., Lelyveld, V.S. and Szostak, J.W. (2018) Inosine, but none of the 8-oxo-purines, is a plausible component of a primordial version of RNA. Proc. Natl. Acad. Sci. U. S. A., 115, 13318–13323.

62. Wright, T.H., Giurgiu, C., Zhang, W., Radakovic, A., O’Flaherty, D.K., Zhou, L.J. and Szostak, J.W. (2019) Prebiotically Plausible “Patching” of RNA Backbone Cleavage through a 3’-5’ Pyrophosphate Linkage. J. Am. Chem. Soc., 141, 18104–18112.

63. Lelyveld, V.S., O’Flaherty, D.K., Zhou, L.J., Izgu, E.C. and Szostak, J.W. (2019) DNA polymerase activity on synthetic N3’-> N5’ phosphoramidate DNA templates. Nucleic Acids Res., 47, 8941–8949.

64. Bhowmik, S. and Krishnamurthy, R. (2019) The role of sugar-backbone heterogeneity and chimeras in the simultaneous emergence of RNA and DNA. Nat. Chem., 11, 1009–1018.

65. Le Vay, K., Weise, L.I., Libicher, K., Mascarenhas, J. and Mutschler, H. (2019) Templated Self-Replication in Biomimetic Systems. Advanced Biosystems, 3.

66. Sosson, M., Pfeffer, D. and Richert, C. (2019) Enzyme-free ligation of dimers and trimers to RNA primers. Nucleic Acids Res., 47, 3836–3845.

67. Zhou, L.J., Kim, S.C., Ho, K.H., O’Flaherty, D.K., Giurgiu, C., Wright, T.H. and Szostak, J.W. (2019) Non-enzymatic primer extension with strand displacement. eLife, 8, 14.

