## Supplemental data for "Deep sequencing of nonenzymatic RNA primer extension"

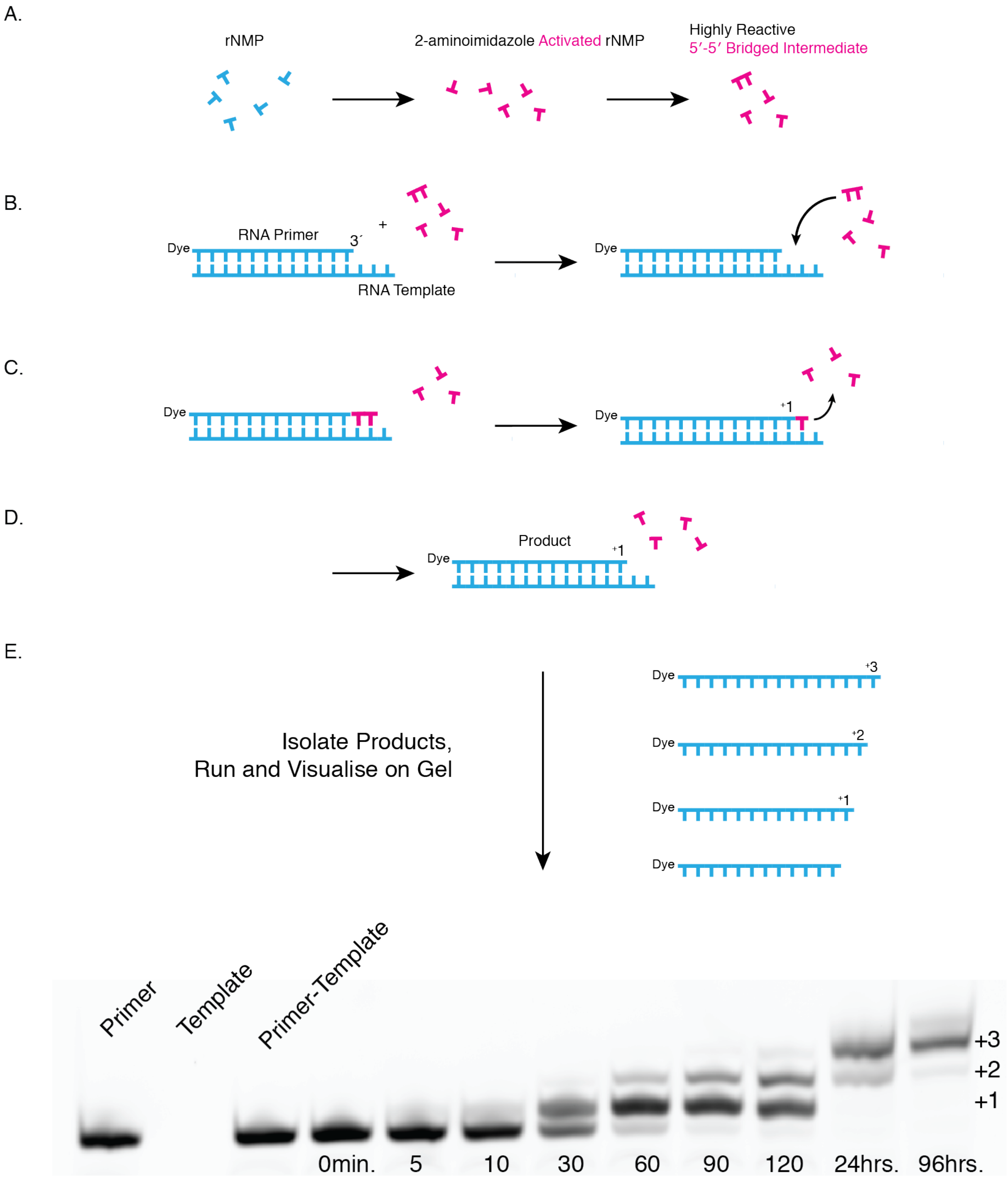


**Figure S1. Schematic of the Mechanism of Nonenzymatic RNA Primer Extension, and an Example of Standard Gel Analysis. A.** Monoribonucleotides are chemically activated with 2-aminoimidazole (2AI) as described in the Material and Methods. Activated mononucleotides spontaneously form a reactive 5´-5´ imidazolium-bridged dinucleotide intermediate in buffered solution. **B.** In the context of an RNA primer-template complex, the dinucleotide intermediate binds and unbinds the template by Watson-Crick base pairing. **C.** In the rate-limiting step, the deprotonated oxygen of the primer 3´ hydroxyl attacks the proximal bridging phosphate, with a 2AI-activated mononucleotide as the leaving group. **D.** The product is a primer extended by one base. **E.** The most common primer extension assay is denaturing (8 M urea)

polyacrylamide gel electrophoresis (PAGE). The product strands, typically labelled with some convenient organic dye, are isolated by the addition of excess competitor that is complementary to the template, and unlabelled. The material is then subjected to denaturing PAGE and the gel is imaged with a fluorescence scanner. The example gel shows the raw data for a typical primer extension time course. (1.2 µM Control Template, 1 µM Control Primer, 200 mM $Na^+$ bicine, pH = 8 and water were heated to 85°C for 30 s, then cooled to 25°C at 0.2 °C/s; 20 mM 2AIrG, 20 mM 2AIrC and 50 mM $MgCl_2$ were added to initiate the reaction. All concentrations indicate final concentrations in a volume of 20 µl. 1 µl aliquots were quenched in 20 µl Urea Load Buffer, 5 µl of which was mixed with 1.3 µl of a 300 µM stock of Control Template Reverse Complement (RC), heated to 95°C for 3 minutes then cooled to 25°C at 0.2 °C/s. 5 µl Urea Load Buffer was added and samples were subjected to denaturing PAGE at 5 W for 20 minutes, then 15 W for 2 hours.)

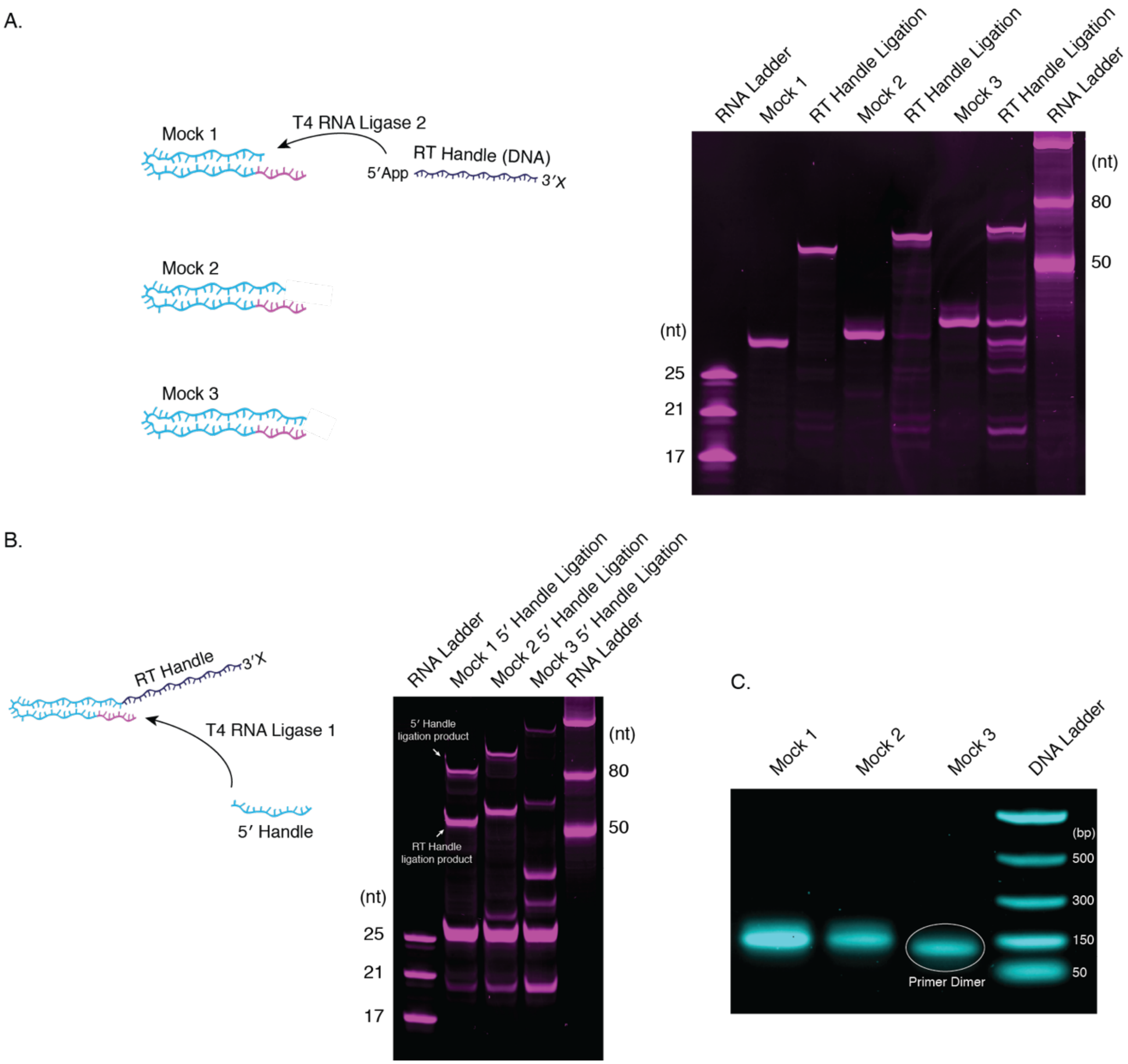


**Figure S2. A Standard RNA-Seq Protocol is Inadequate for Processing the Hairpin Construct. A.** Three mock hairpin constructs (Mock 1, Mock 2 and Mock 3; 31 nt, 34 nt and 38 nt, respectively) representing different apparent extents of primer extension (0, +3 and +7, respectively) were ligated to the RT Handle (21 nt). Reactions were performed with 3.6 µM mock substrate, 5.4 µM RT Handle 1 and NEBNext® 3´ Ligation Enzyme Mix, and incubated for 18 hours at 16°C. Mock 1 and Mock 2 ligate well, but Mock 3 does not, with a significant residual unligated band as well as off-target products (target length = 59 nt; off-target bands appear to run faster). **B.** The products from the RT handle ligation were then ligated to the 5´ Handle. Reactions were performed with ~ 2.7 µM substrate, 4 µM 5´ Handle, 7% $PEG_{8000}$ and 30 U of T4 RNA Ligase 1, and incubated for 1 hour at 16°C. The ligation efficiencies are low for all constructs, and particularly low for Mock 3. Off-target products are again prominent in the case of Mock 3. **C.** The products of the 5´ ligation reaction were used for barcoding PCR, and the PCR products were analysed by agarose gel electrophoresis. Mock 1 and Mock 2 yield the expected products (150 bp and 153 bp), but Mock 3 yields only primer dimer (127 bp).

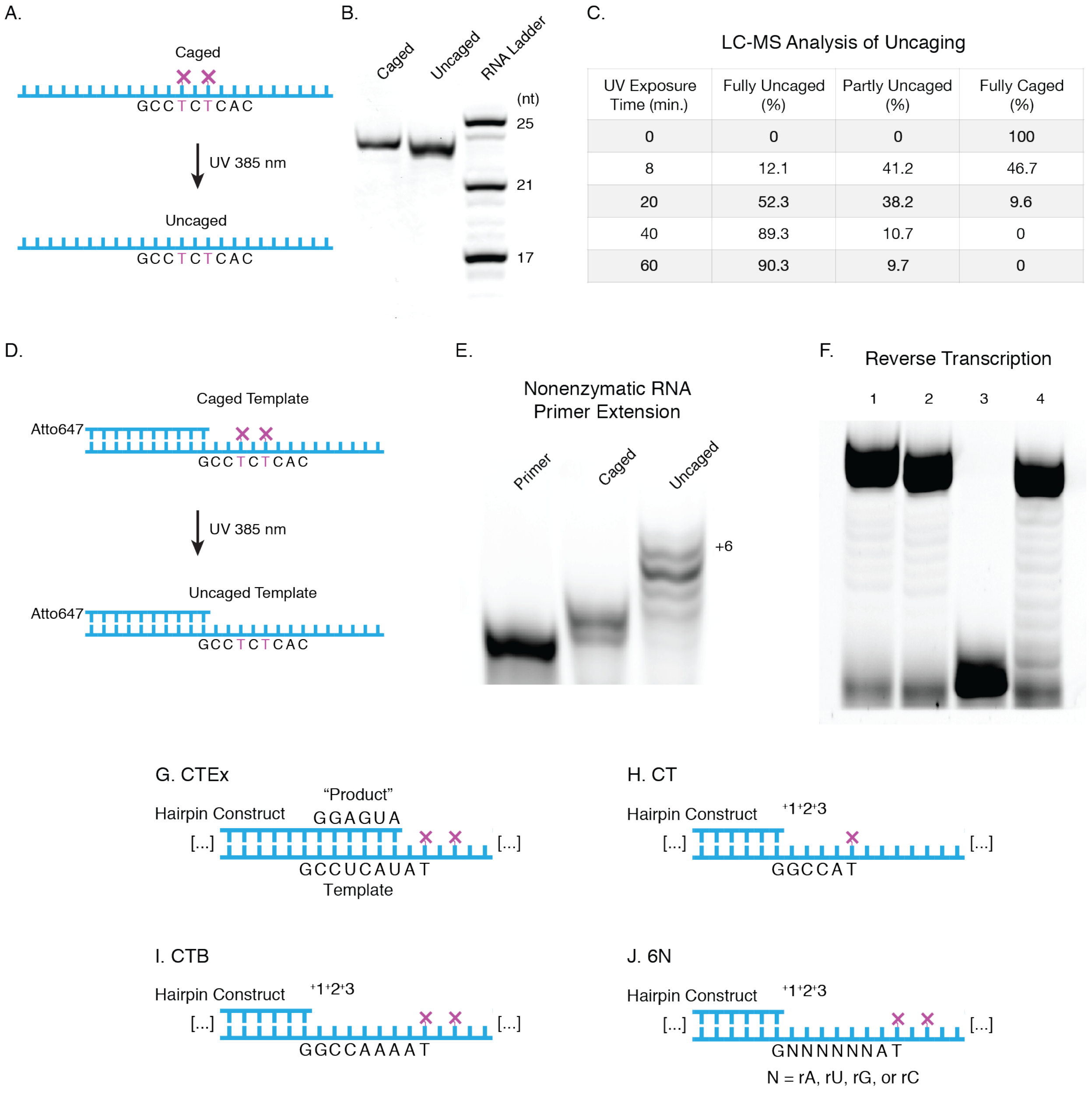


| UV Exposure Time (min.) | Fully Uncaged (%) | Partly Uncaged (%) | Fully Caged (%) |
|---|---|---|---|
| 0 | 0 | 0 | 100 |
| 8 | 12.1 | 41.2 | 46.7 |
| 20 | 52.3 | 38.2 | 9.6 |
| 40 | 89.3 | 10.7 | 0 |
| 60 | 90.3 | 9.7 | 0 |

**Figure S3. Caged Bases are Effective at Halting Primer Extension and Reverse Transcription. A.** The RNA NPOM Test Oligo harbours two NPOM-caged dT bases. **B.** When 1 µM of the oligo in 10 µl of 10 mM Tris-Cl (pH = 8) is irradiated with 385 nm UV for 8 minutes, the uncaging registers as a gel shift by PAGE analysis. (2.5 µl of each sample was added to 7.5 µl Urea Load Buffer and subjected to denaturing 18% PAGE at 18 W for 2 hours.) **C.** LC-MS was then used to identify optimal conditions for full uncaging (10 µM samples in 15 µl water), with no detectable fully caged and almost 90% fully uncaged oligo after 40 minutes of irradiation. **D-E.** Primer extension is completely stalled by a caged template, whereas the uncaged template functions normally. Primer extension was performed in a final volume of 20 µl with 1 µM primer, 1.2 µM template, 200 mM $Na^+$ bicine, pH = 8, 20 mM $MgCl_2$ and 20 mM each of 2AIrG and 2AIrA. The template and an appropriate volume of water were UV irradiated, and the primer added. The mixture was then heated to 85°C for 30 s and cooled to 25°C at 0.2 °C/s to anneal the primer before other ingredients were added. The reaction was quenched by the addition of 1 µl of the reaction mixture to 19 µl Urea Load Buffer. 5 µl of the quenched mixture was added to 1.3 µl of a 300 µM stock of NPOM Test Oligo RC, which was then heated to 95°C for 3 minutes and cooled to 25°C at 0.2 °C/s. 5 µl of Urea Load Buffer was added and samples were subjected to denaturing PAGE at 5 W for 15 minutes then 15 W for 1 hour. **F.** A DNA primer was used in place of the RNA primer, and the template was tested for reverse transcription (RT). 1: RT on a control template with no caged bases. 2: RT on a control template with no caged bases after irradiation with 385 nm UV. 3: RT on the NPOM Test Oligo template. 4: RT on the uncaged NPOM Test Oligo template. RT reactions were performed in a final volume of 30 µl with 2.7 µM template, 3 µM primer, 8 µl First Strand Synthesis Buffer (NEB), 1 µl Murine RNase Inhibitor (NEB) and 1 µl ProtoScript® II at 50°C for 1 hour. PAGE analysis was as in (D), except 4.4 µl of the reaction mixture was quenched in 15.6 µl Urea Load Buffer and 0.66 µl of the quenched mixture was added to the NPOM Test Oligo RC. **G-J.** The RNA hairpin constructs used in this study all harbour the same essential features: a defined hairpin, a priming rC base, a variable template region, a placeholding rA at the end of the template, NPOM-caged dT bases and an already-encoded downstream 5´ Handle. **G.** The Control Template Extended construct (CTEx) has a defined template sequence 3´-CCUCAU-5´ and a defined product sequence 5´-GGAGUA-3´, mimicking perfect and complete primer extension. **H.** The Control Template construct (CT) was the first version of a defined-template-sequence test hairpin, and was replaced by **I.,** Control Template B (CTB), which more closely resembles the ultimate experimental construct (J). CT and CTB both harbour a 3´-GCC-5´ template sequence. **J.** The 6N construct harbours a template with six randomized positions, representing all possible templating 6-mers.

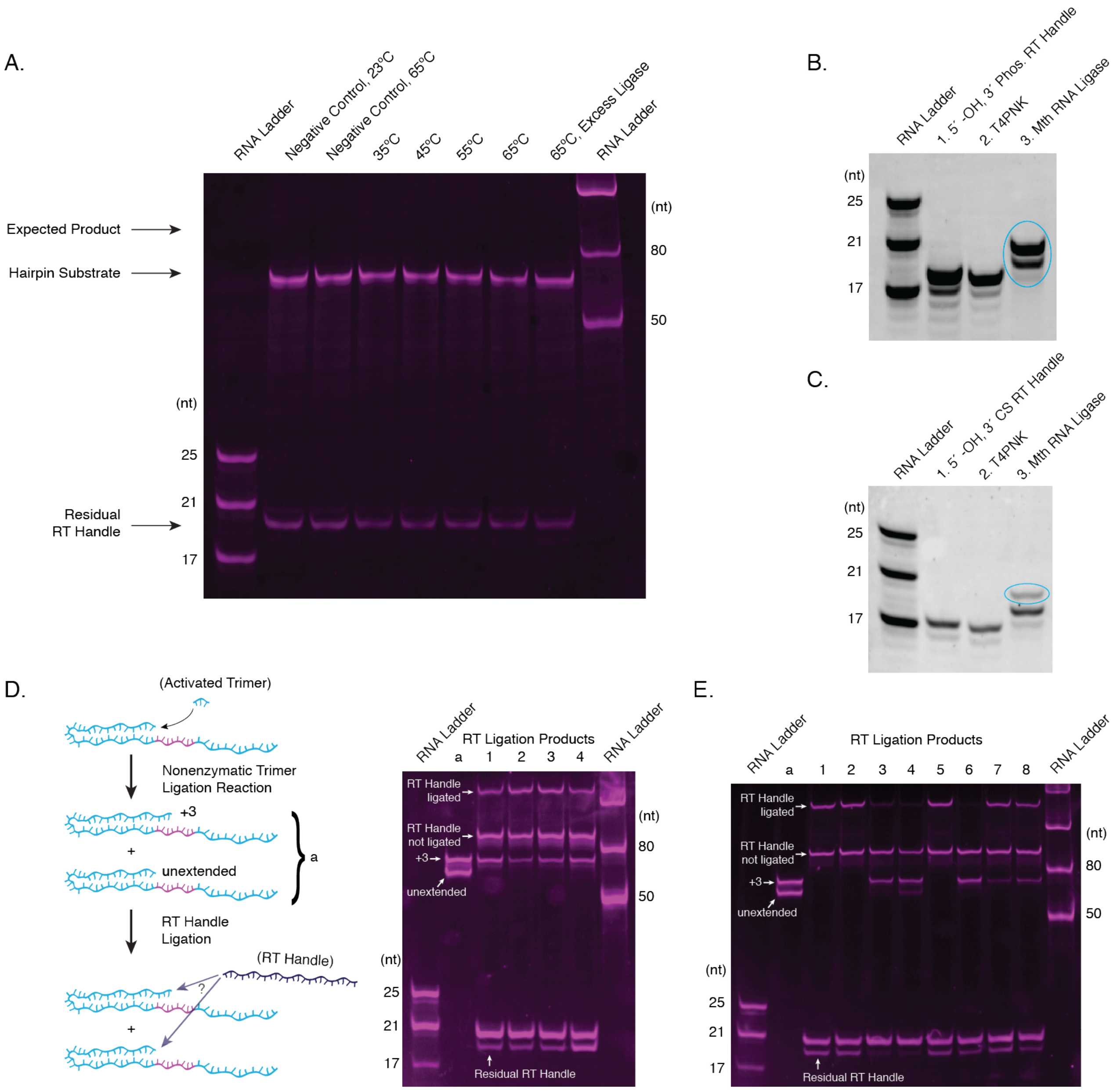


**Figure S4. Optimising RT Handle Ligation. A.** Thermostable App Ligase is not active for RT Handle ligation to the hairpin construct. Reaction conditions were 1 µM 6N (59 nt), 1.2 µM RT Handle 2 and 2.6 µM Thermostable App Ligase (NEB) incubated for 1 hour at the indicated temperatures. The Negative Control sample did not include enzyme, and the Excess Ligase sample had 5.2 µM enzyme. 1 µl of each sample was quenched in 20 µl Urea Load Buffer and subjected to denaturing PAGE. **B-C.** Regardless of blocking chemistry or vendor source, pre-adenylated RT handles exhibit an impurity, visible by PAGE analysis (see for example "Residual RT Handle" in the gels shown in (D) and (E), where the RT Handle registers as two bands rather than one). Vendor-supplied material was not concentrated enough to gel purify the target band, so we attempted several in-house preparations. **B.** We synthesized an RT Handle sequence with a free 5´ (-OH) and an ostensibly blocking 3´ phosphate. Following phosphorylation with T4 Polynucleotide Kinase (3´ phosphatase minus) (T4PNK), we found that the adenylating enzyme Mth RNA Ligase recognizes both 3´ and 5´ phosphates, visible as a higher band with two adenylations, and a lower band with one adenylation. Such products could severely interfere with the target ligation reaction.

The phosphorylation reaction was 6 µM (5´-OH, 3´-phosphate RT Handle) and 10 U T4PNK in a final volume of 50 µl, incubated for 30 minutes at 37°C. Samples were purified with a Zymo Research Oligo Clean & Clear Concentrator™ spin column, eluted in 30 µl water and 0.1 µl in 20 µl of Urea Load Buffer was used for PAGE. 9 µl of the eluate was adenylated in a reaction with 100 pmole Mth RNA Ligase in a final volume of 20 µl. Samples were purified with a Zymo Research Oligo Clean & Clear Concentrator™ spin column, eluted in 25µl water and 0.25 µl in 20 µl of Urea Load Buffer was used for PAGE. **C.** We next tested a 3´ three-carbon-spacer (3CS) block using the same protocol as in B., but found that T4PNK appears to phosphorylate the 3´ end as well, resulting, again, in some 3´ adenylation. We interpret this to mean that T4PNK can recognise the terminal -OH on the 3CS, and that Mth RNA ligase can recognise a phosphate attached to a 3CS. Ultimately, the final optimised protocol employs a 3´ dideoxy, 5´ pre-adenylated oligo. The persistent impurity, most likely unadenylated material, does not appear to interfere with ligation or the results in any measurable way (see Figures 2A and S7). **D-E.** RT Handle ligation tests on nonenzymatic ligation products. The +3 product of a nonenzymatic ligation reaction on CTB with the activated 2AI-CGG trimer proved inefficient in RT Handle ligation reactions. **D.** The product of a nonenzymatic ligation reaction (CTB + 2AI-CGG, (a)) yields a clearly discernible +3 product band and an unextended band by PAGE. Unextended material can then be fully ligated to RT Handle, but the +3 product is only partially ligated (see residual +3 band under condition 1, for example). Note that the hairpin construct can run in unexpected ways during PAGE after RT Handle ligation: the product of RT Handle ligation to the unextended material runs significantly faster than the product of RT Handle ligation to the +3 product, despite only a three base difference (see also Figures 2, S7 and S8). CTB nonenzymatic ligation reactions were performed as per the standard NERPE protocol, but without the 5´ Handle Block. All samples were heated to 95°C for 2 minutes and cooled to 70°C at 1 °C/s, followed by the addition of RT Handle such that the final concentration in the reaction would equal 2 µM. The mixture was then heated to 70°C for 2 minutes and cooled to 23°C at 1 °C/s. Reactions were in a final volume of ~14 µl with 12% $PEG_{8000}$ and 3 µM T4 RNA Ligase 2, truncated KQ, and incubated for 24 hours at 16°C unless otherwise noted. 1: Standard RT Handle ligation reaction, as described above. 2: Reaction supplemented with ~10% DMSO. 3: Reaction incubated at 25°C. 4: 6 µM ligase and reaction incubated at 25°C. The addition of DMSO and a 25°C incubation slightly improved the reaction. **E.** Reactions were as in D., but the incubation was for 18 hours at 25°C, unless otherwise noted, followed by two hours at 4°C. 1: Reaction supplemented with ~10% DMSO. 2: Reaction supplemented with ~20% DMSO. 3: Reaction supplemented with ~10% DMSO and incubated at 37°C. 4: Reaction supplemented with ~20% DMSO and incubated at 37°C. 5: Reaction supplemented with 20% $PEG_{8000}$. 6: $PEG_{8000}$ omitted. 7: Reaction supplemented with 1 mM hexamminecobalt chloride (1). 8: ~ 1.5 µM ligase. DMSO combined with a 25°C incubation, (1) and (2), or excess $PEG_{8000}$, (5), were equally effective in driving the RT Handle ligation reaction to completion.

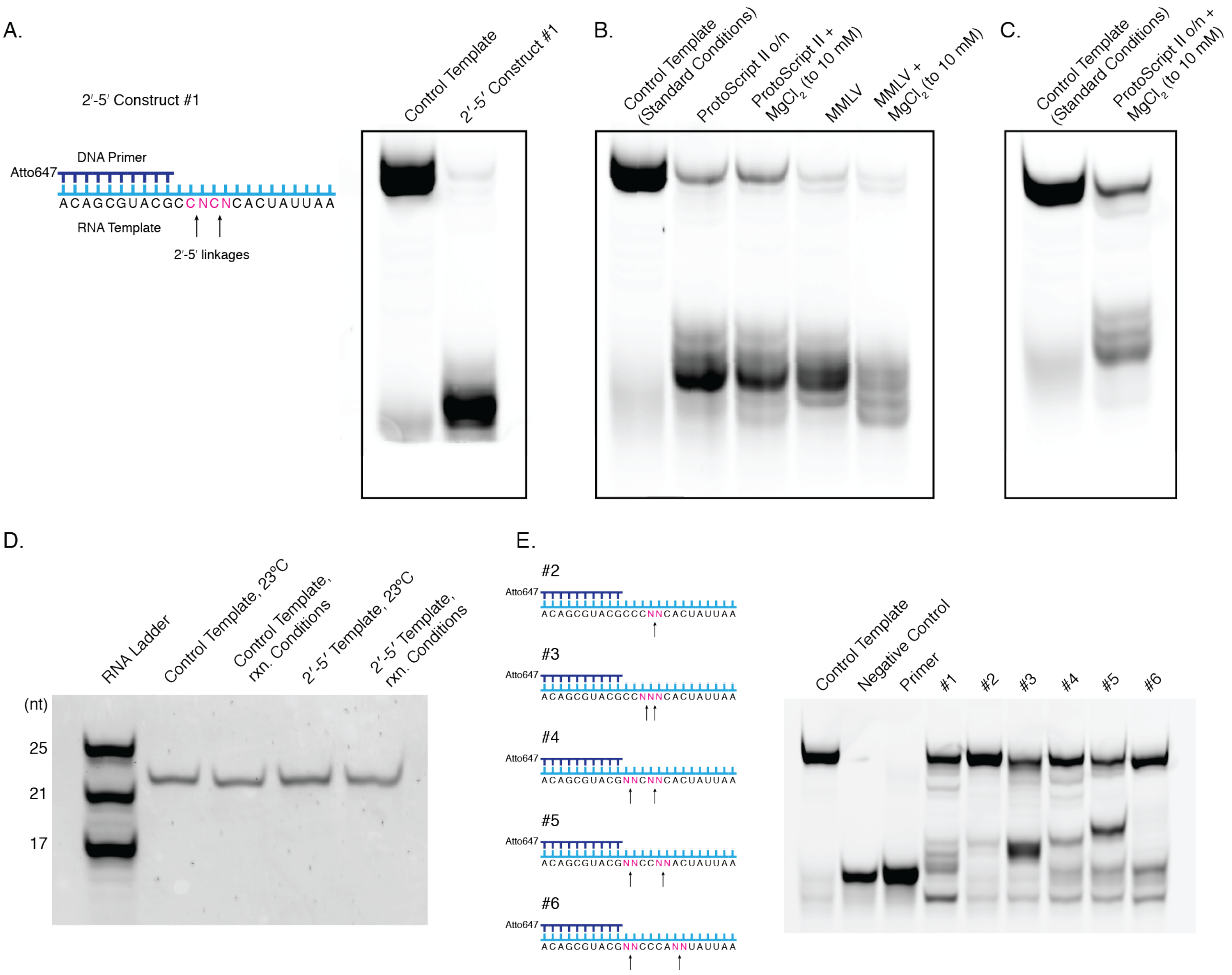


**Figure S5. The Effect of 2´-5´ Linkages on Reverse Transcriptase Activity. A.** A reverse transcriptase test construct (2´-5´ Construct #1) with two non-consecutive 2´-5´ linkages (arrows), each with a randomized downstream base. ProtoScript® II reverse transcriptase yields the expected full-length DNA product on a control template with no 2´-5´ linkages, but is almost completely stalled by the 2´-5´ linkages in the test construct. Reactions were in a final volume of 40 µl with 2.7 µM template and 3 µM primer. The template and primer were heated in water to 85°C for 30 s, then cooled to 25°C at 0.2 °C/s. 8 µl NEBNext® First Strand Synthesis Buffer (NEB), 1 µl Murine RNase Inhibitor (NEB) and 1 µl ProtoScript® II were added to the mixture and incubated at 50°C for 60 minutes. The reaction was quenched by aliquoting 4.4 µl into 15.6 µl Urea Load Buffer. A 0.66 µl aliquot of the quenched reaction was mixed with 1.3 µl of a 300 µM stock of RT Test RC and 5 µl Urea Load Buffer, heated to 95°C for 3 minutes, then cooled to 25°C at 0.2 °C/s. 5 µl more Urea Load Buffer was added and samples were subjected to denaturing PAGE at 5 W for 15 minutes, then 15 W for 1 hour. **B.** Screen of conditions on the same construct, with the goal of increasing full-length product yield. Reactions were prepared as in (A), except with 2 µl ProtoScript® II or MMLV reverse transcriptase, no Murine RNase Inhibitor, MMLV reaction buffer in MMLV reactions and reverse transcription was at 42°C for 3 hours. The Control Template (Standard Conditions) is identical to that used in (A). o/n indicates overnight = 12 hours. A longer incubation time and increasing the concentration of $MgCl_2$ to 10 mM both increase yields. **C.** Combining the 12-hour incubation time with $MgCl_2$ to 10 mM yields the best results from among conditions tested. Reactions were prepared as in (B), but with 50 mM Tris-Cl (pH = 8), 75 mM KCl, 10 mM DTT, 10 mM $MgCl_2$, 1 mM each dNTP and 800 U ProtoScript® II. The Control Template (Standard Conditions) is identical to that used in (A). o/n indicates overnight = 12 hours. **D.** The conditions identified in (C) do not degrade the RNA template. Templates were prepared as in (C) but without enzyme or primer, and 1.5 µl of of the quenched reaction was mixed with 15 µl Urea Load Buffer, heated to 95°C for 1 minute, then cooled to 25°C at 1 °C/s. Samples were subjected to denaturing PAGE at 5 W for 10 minutes then 15 W for 1 hour and stained with SYBR Gold™ for imaging. rxn. Conditions indicate reverse transcription reaction conditions equivalent to that used in (C), and the 2´-5´ Template is the same as used in 2´-5´ Construct #6 (E). **E.** A series of constructs, including the Control Template, were tested under the reverse transcription conditions identified in (C). Reactions were performed as in (C). Samples were quenched by aliquoting 4.4 µl into 4.6 µl water with 1 µl of 1 M NaOH. The mixture was heated to 90°C for 10 minutes and neutralized with 1 µl of 1 M HCl (2) before the addition of 13.4 µl Urea Load Buffer. 0.6 µl of of the quenched mixture was added to 15 µl Urea Load Buffer, heated to 95°C for 1 minute, then cooled to 25°C at 1 °C/s. Samples were subjected to denaturing PAGE at 5 W for 10 minutes then 15 W for 1 hour. The Negative Control included all ingredients except the enzyme, and the Primer lane included only primer with buffer. All lanes derived from reactions that included enzyme show a band of unknown origin down-shifted from the primer band (compare for example "Primer" and "#1").

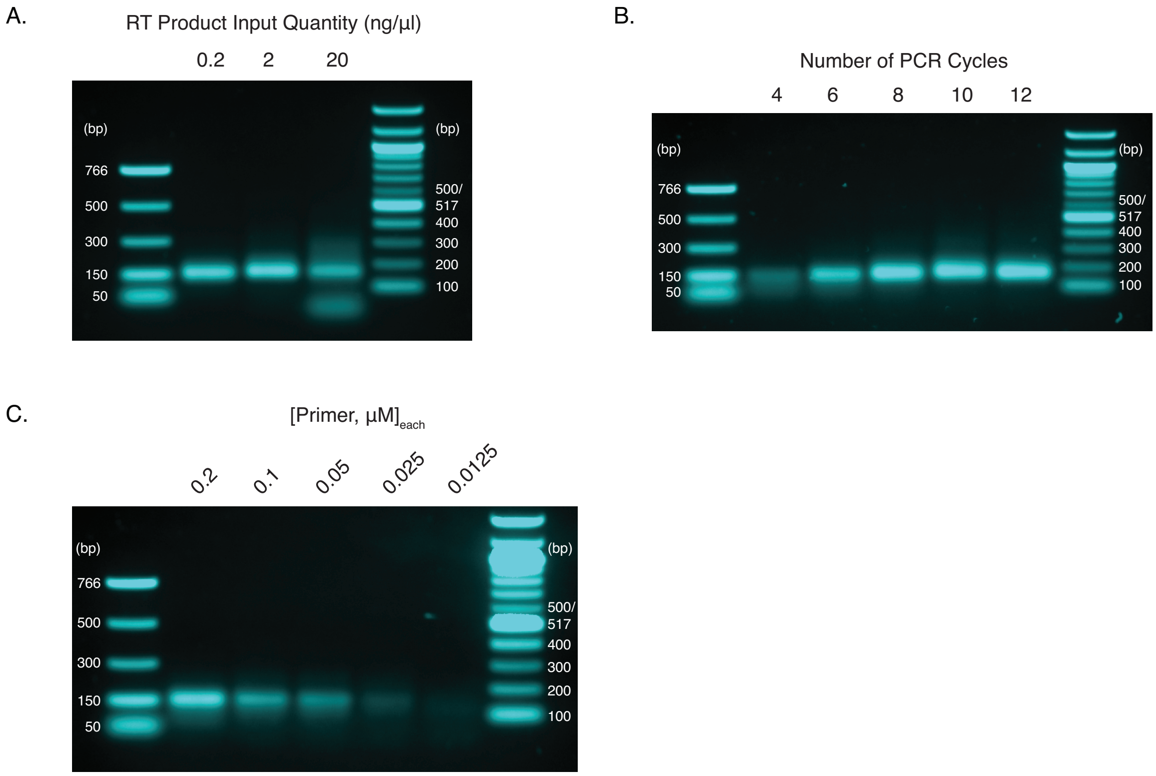


**Figure S6. Optimising PCR Conditions.** PCR reactions and non-preparative agarose gels were prepared as described in the Material and Methods. The product of RT Handle ligation to the 6N construct was used as a test PCR template. **A.** PCR products across a titration of input cDNA quantities. The protocol benefits from maximizing the quantity of input material. 2 ng/µl is the highest tested quantity that does not result in side products. The PCR reaction was with 12 cycles and 0.2 µM of each primer. **B.** PCR products across increasing numbers of PCR cycles. The protocol benefits from minimising the number of PCR cycles, and 6 cycles yielded a robust enough band for downstream usage. The PCR reaction was with 2 ng/µl of template, 12 cycles, and 0.2 µM of each primer. **C.** PCR products across a range of PCR primer concentrations. The protocol benefits from minimising the concentrations of the primers, and we reasoned that if there were sufficient quantities of the primers after PCR with the conditions identified in (A-B), then a reduction in the primer concentration would not reduce the quantity of product. However, even the change from 0.2 µM of each primer to 0.1 µM of each primer reduces product yield, suggesting that the primer concentrations should not be reduced below 0.2 µM.

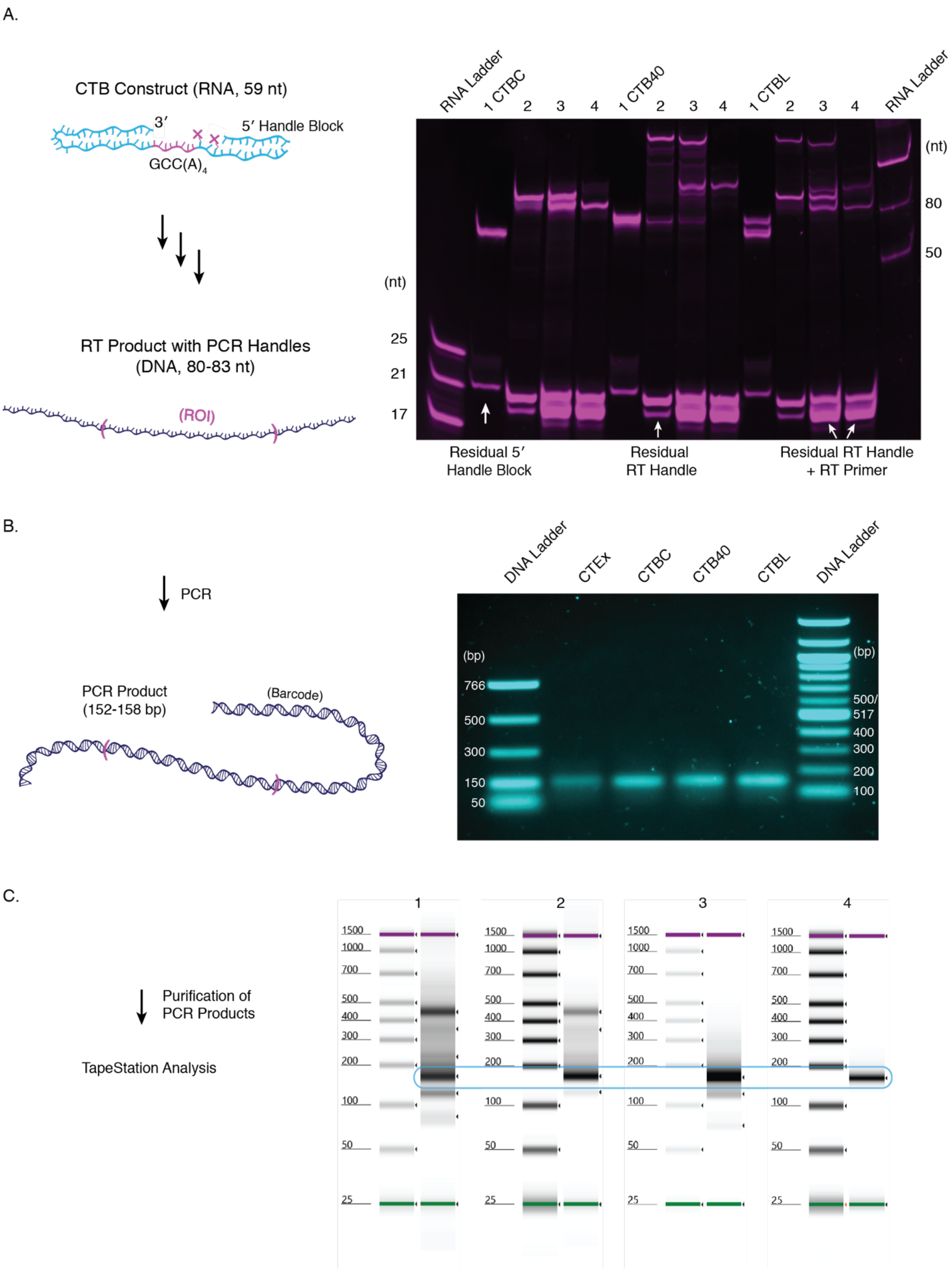

A.
CTB Construct (RNA, 59 nt)
3′
5′ Handle Block
GCC(A)4
RT Product with PCR Handles
(DNA, 80-83 nt)
(ROI)
RNA Ladder
1 CTBC
2
3
4
1 CTB40
2
3
4
1 CTBL
2
3
4
RNA Ladder
(nt)
80
50
(nt)
25
21
17
Residual 5′ Handle Block
Residual RT Handle
Residual RT Handle + RT Primer
B.
PCR
PCR Product
(152-158 bp)
(Barcode)
DNA Ladder
CTEx
CTBC
CTB40
CTBL
DNA Ladder
(bp)
766
500
300
150
50
(bp)
500/
517
400
300
200
100
C.
Purification of PCR Products
TapeStation Analysis
1
2
3
4
1500
1000
700
500
400
300
200
100
50
25

**Figure S7. Conversion of Nonenzymatic RNA Primer Extension Products to dsDNA Ready for Sequencing. A.** The products of control experiments using the CTB construct were converted into cDNA using the optimised protocol. CTB Control (CTBC) = mock NERPE; CTB, 40 mM total monomer (CTB40) = NERPE with 20 mM 2AIrG and 20 mM 2AIrC; CTB Ligation (CTBL) = nonenzymatic ligation with 500 µM 2AI-CGG. For each experiment 1 = post desalting, 2 = RT Handle ligation, 3 = reverse transcription, and 4 = post cDNA isolation. Compare CTBC with Figure 2A. Note that banding patterns for CTB40 and CTBL are from the multiple products of NERPE (for example, unextended and + 3 products for CTBL). Note also that in the case of CTB40 the residual band of unligated material after RT Handle ligation (CTB40, 2) is not specific to any particular product because the distribution of NERPE products as analysed by PAGE and NERPE-Seq agree (see Figure 4); we conclude that the residual band resulted from incomplete ligation of the RT Handle overall rather than a bias against any specific NERPE product. **B.** The optimised PCR conditions (Figure S6) yield bands of the expected sizes across constructs. Samples were purified with a QIAGEN QIAquick® PCR Purification spin column prior to agarose gel analysis. **C.** TapeStation analysis of the PCR products is more sensitive than agarose gel imaging and can identify unwanted side-products. PCR reactions were purified using 1: QIAGEN QIAquick® PCR Purification spin column, 2: Agencourt® AMPure® beads, 3: Quantum Prep® Freeze 'N Squeeze agarose gel purification spin columns or 4: both Agencourt® AMPure® beads and Quantum Prep® Freeze 'N Squeeze agarose gel purification spin columns. 4 yields the best results, with a well-defined band devoid of any detectable side-products. Note that the PCR reaction had not yet been optimised at 1.

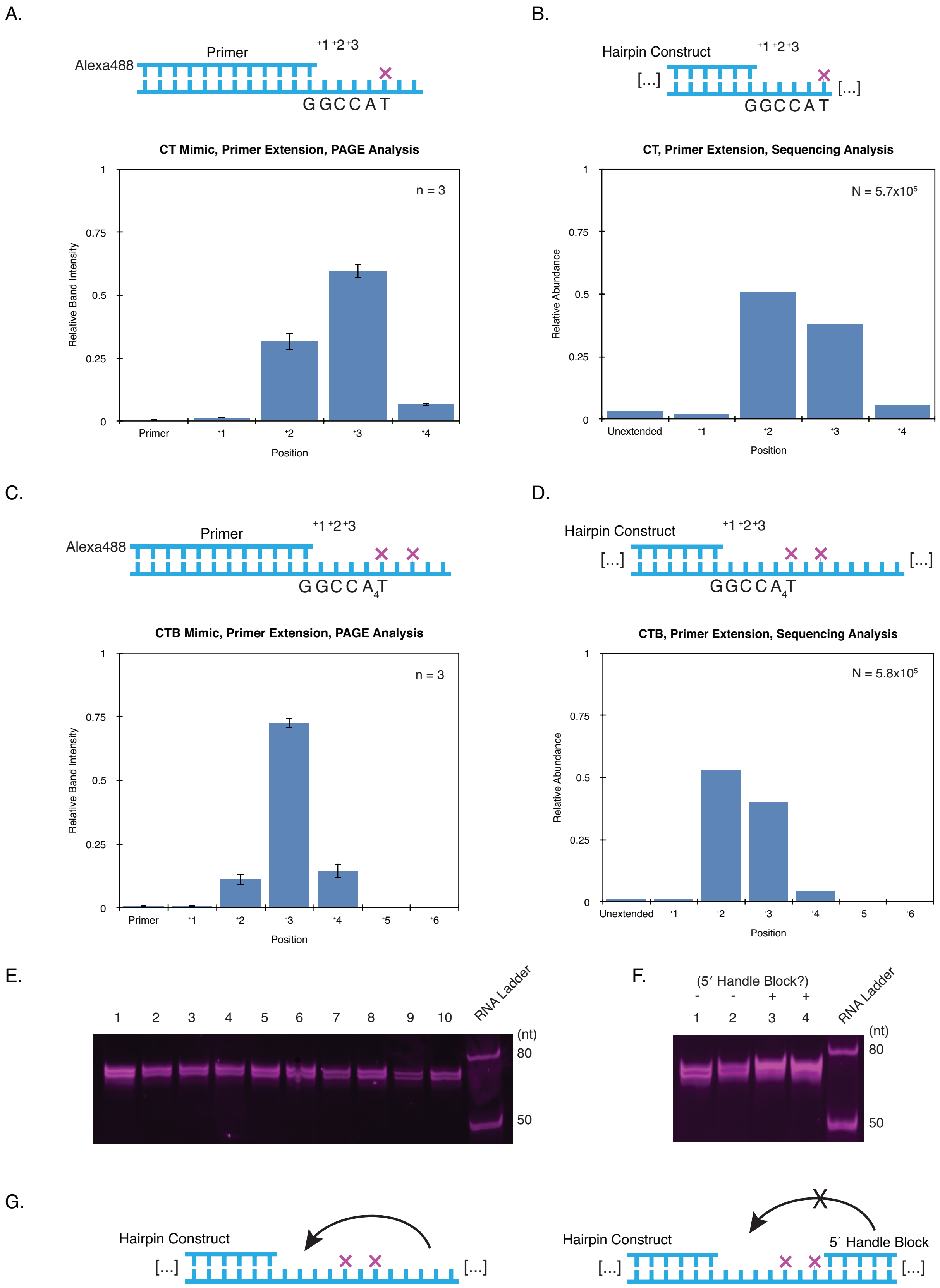
A.
Primer
Alexa488
+1 +2 +3
GGCCAT
CT Mimic, Primer Extension, PAGE Analysis
n = 3
Relative Band Intensity
1
0.75
0.5
0.25
0
Primer
+1
+2
+3
+4
Position
B.
Hairpin Construct
[...]
[...]
+1 +2 +3
GGCCAT
CT, Primer Extension, Sequencing Analysis
N = 5.7x10^5
Relative Abundance
Unextended
+1
+2
+3
+4
Position
C.
Primer
Alexa488
+1 +2 +3
GGCCA4T
CTB Mimic, Primer Extension, PAGE Analysis
n = 3
Relative Band Intensity
Primer
+1
+2
+3
+4
+5
+6
Position
D.
Hairpin Construct
[...]
[...]
+1 +2 +3
GGCCA4T
CTB, Primer Extension, Sequencing Analysis
N = 5.8x10^5
Relative Abundance
Unextended
+1
+2
+3
+4
+5
+6
Position
E.
1 2 3 4 5 6 7 8 9 10
RNA Ladder
(nt)
80
50
F.
(5′ Handle Block?)
- - + +
1 2 3 4
RNA Ladder
(nt)
80
50
G.
Hairpin Construct
[...]
[...]
Hairpin Construct
5′ Handle Block
[...]
[...]

**Figure S8. The 5´ Handle Interferes with Primer Extension. A.** PAGE analysis of the length distribution of primer extension products using a template meant to mimic the Control Template (CT) hairpin construct (B). Reaction conditions were as in Figure S1E, and the reaction time was 24 hours. **B.** NERPE-Seq analysis of the same reaction as in (A), but using the CT construct. **C.** PAGE analysis of the length distribution of primer extension products using a template meant to mimic the CTB hairpin construct (D). **D.** NERPE-Seq analysis of the same reaction as in (A), but using the CTB construct. **E.** PAGE of primer extension products using CTB across hairpin folding conditions. The expected result of this primer extension is primarily +3 products, as in (C). However, primer extension on CTB yields significant quantities of both +2 and +3 products (D), and these two prominent bands can be visualised, though not accurately quantified, by PAGE analysis. 1: Standard NERPE-Seq reaction conditions as described in the Material and Methods, without the 5´ Handle Block, 2: water to adjust for the final volume added after annealing, 3: 1 mM NaCl supplemented to annealing step, 4: annealing final temperature set to 4ºC, 5: temperature ramp down at 0.1 ºC/s, 6: heating for 6 minutes at 98ºC, 7: heating for 6 minutes at 98ºC, temperature ramp down at 0.1 ºC/s, and annealing final temperature set to 4ºC, 8: primer extension performed at 18ºC, 9: activated monomers not pre-mixed, 2AIrG added first, 10: activated monomers not pre-mixed, 2AIrC added first. The reaction products were desalted and subjected to denaturing PAGE at 5 W for 10 minutes then 23 W for 1 hour and 10 minutes and stained with SYBR Gold™ for imaging. The proportions of +2 and +3 products are the same in all cases, eliminating the tested variables as potential causes for the discrepancy between (C) and (D). **F.** The assay was the same as in (E). 1 and 2 were the same reaction as in (E, lane 1), and 3 and 4 included 1.2 µM 5´ Handle Block Test during the primer extension reaction. 3 and 4 reproduce the expected PAGE signature of primarily +3 products. 1: Standard reaction conditions, 2: standard reaction conditions with 5´ Handle Block Test added immediately prior to gel loading, 3: 1.2 µM 5´ Handle Block Test included during the primer extension reaction, products not heated in Urea Load Buffer prior to gel loading, 4: 1.2 µM 5´ Handle Block Test included during the primer extension reaction, products heated in Urea Load Buffer at 95ºC for 3 minutes prior to gel loading. **G.** When the 5´ Handle of CT or CTB is single-stranded it can transiently interact with the primer extension templating sequence and inhibit the reaction. This accounts for the discrepancies between (A) and (B), and (C) and (D). But if the 5´ Handle is occupied by a complementary oligo, it becomes inert and does not affect primer extension. See also Figure 4.

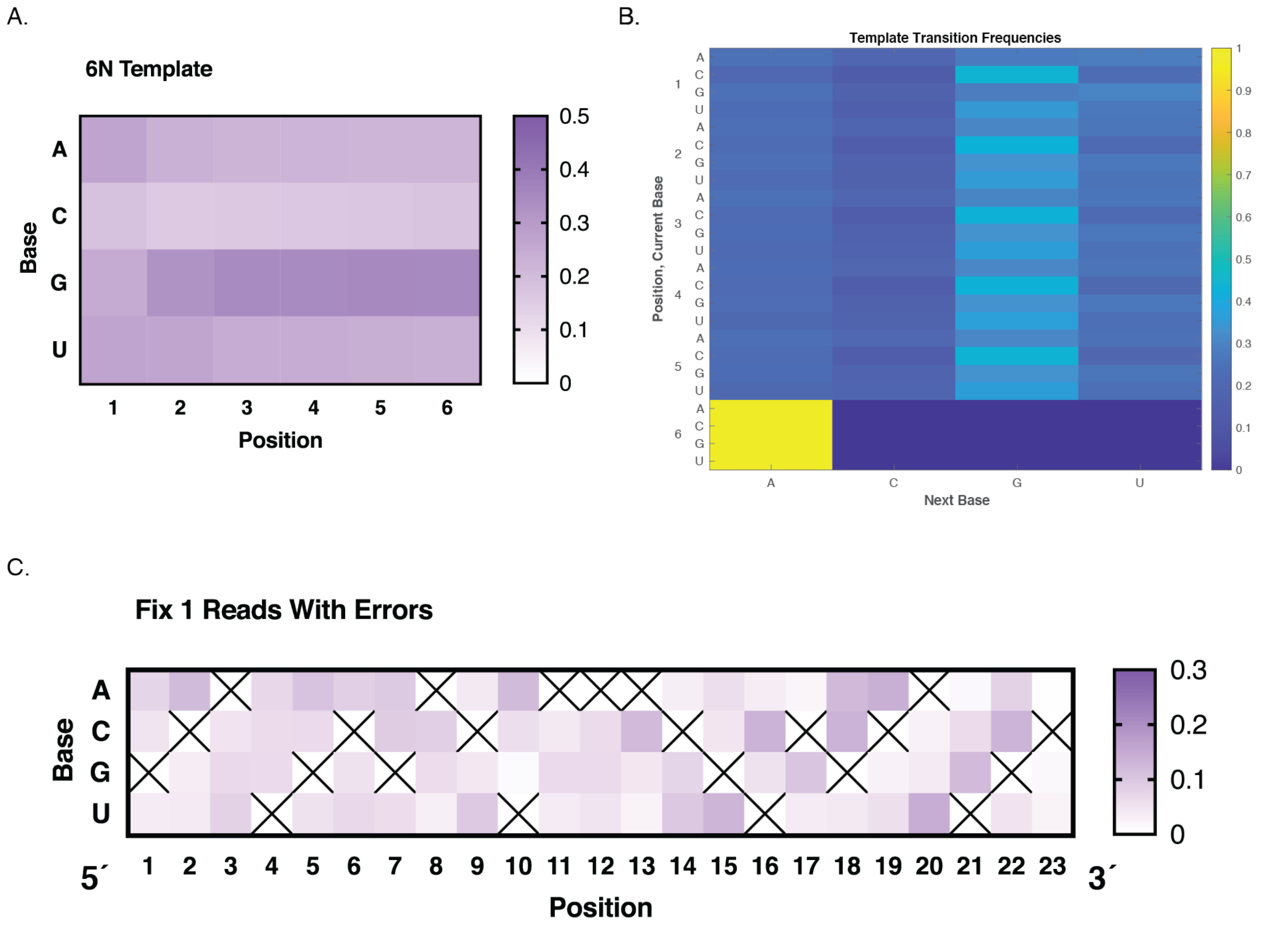


**Figure S9. Solid-state Oligonucleotide Synthesiser Biases and Errors. A.** The position-dependent base frequencies of the 6N template. The template sequence was synthesized using an equimolar mix of all four phosphoramidites. An ideal distribution would be 0.25 of each base at each position, but there are clear deviations, highlighting the need for normalising data during analysis where possible. In particular, the distribution becomes progressively skewed towards Gs and away from Cs. **B.** The position-dependent base transition frequencies of the 6N template. Ideally, all transitions would occur with equal probabilities of 0.25, but the heatmap shows structure. Note the consistent high frequency of transitions to G (especially C-to-G) and the low frequency of transitions to C (especially C-to-C and U-to-C). This accounts for the distribution in (A). Transitions *to* A and *from* G or A are more uniform. The analysis included the final placeholding A in position 7 as an internal control, so all final transitions *from* position 6 are "to A". **C.** The position-dependent base frequencies of the Fix 1 sequence (which encodes the hairpin) in the 6N construct from forward reads that contained at least one error. Even reads with at least one error are mostly correct, so the frequency of the correct base at each position is high. To show the frequencies of the incorrect bases, the scale was adjusted down and an X marks the correct base at each position. The RNA hairpin constructs were synthesised in the 3´-to-5´ direction. Error frequencies vary, but we note that in over half of positions the highest frequency incorrect base is the same as the preceding correct base in the synthesis direction. This could simply result from contamination by the previously added base, or the addition of two bases in one step. The trend is more pronounced toward the 3´ end, though it is unclear why the artefact should become less common as a function of the number of synthesis cycles. Oligonucleotide synthesis is performed in organic solvent, and the phosphoramidites carry a variety of protecting groups, some of which are unique to specific bases. Therefore, the traditional biochemical view of base stacking interactions in aqueous buffer, for example, may not be relevant. Nonetheless, this result indicates that there are non-trivial interactions between already-incorporated nucleotides and incoming phosphoramidites. These could arise from specific affinities between chemical groups, or from different degrees of steric hindrance among specific pairwise combinations that could impact reaction rates. The solution is not so simple as the base-dependent occlusion of phosphoramidite reactive groups because then all pair-wise combinations involving a given base would be exhibit the same pattern. Similarly, the fact that the biases are not symmetric (G-to-C is not favored, for example), indicates that the chemical status of the already-incorporated base is key.

### Table of Oligonucleotides

| Oligo Name | Type | Source | Sequence (5′-3′; all termini free -OH unless otherwise noted) | Notes |
|---|---|---|---|---|
| Control Primer | RNA | IDT† | Alexa488-AGUGAGUAACUC | Figure S1 |
| Control Template | RNA | IDT | CCGGAGUUACUCACU | Figure S1 |
| Control Template RC‡ | RNA | IDT | AGUGAGUAACUCCGG | Figure S1 |
| Mock 1 | RNA | IDT | Phos.-AAAACCCCGCAUGCGACUAAACGUCGCAUGC | Figure S2, standard desalting purification only |
| Mock 2 | RNA | IDT | Phos.-AAAACCCCGCAUGCGACUAAACGUCGCAUGCGGG | Figure S2, standard desalting purification only |
| Mock 3 | RNA | IDT | Phos.-AAAACCCCGCAUGCGACUAAACGUCGCAUGCGGGUUU | Figure S2, standard desalting purification only |
| RT Handle 1 | DNA | NEB | App-AGATCGGAAGAGCACACGTCT-$NH_2$ | Figure S2 |
| 5′ Handle (SR Adaptor) | RNA | NEB or IDT | GUUCAGAGUUCUACAGUCCGACGAUC | Figure S2 |
| NPOM Test Primer | RNA | IDT | Atto647-UGUCGCAUGC | Figure S3, for NERPE |
| RT Test Primer | DNA | IDT | Atto647-TGTCGCATGC | Figures S3 and S5 |
| NPOM Test Oligo | RNA/DNA | In-house⊥ | AAUUAUCACdT(-NPOM)CdT(-NPOM)CCGCAUCCGACA | Figure S3 |
| RT Control Template | RNA | IDT | AAUUAUCACCCCCCGCAUGCGACA | Figures S3 and S5 |
| NPOM Test Oligo RC | RNA | IDT | UGUCGCAUGCGGGGGGUGAUAAUU | Figure S3 |
| RT Handle 2 | DNA | IDT | App-AGATCGGAAGAGCACACGTCT-3CS | Figure S4 |
| 5′-OH, 3′-phos. RT Handle | DNA | In-house | AGATCGGAAGAGCACACGTCT-Phos. | Figure S4 |
| 5′-OH, 3′-CS RT Handle | DNA | In-house | AGATCGGAAGAGCACACGTCT-3CS | Figure S4 |
| 2′-5′ Construct #1 | RNA | In-house | AAUUAUCACN*CN*CCGCAUGCGACA | Figure S5 |
| 2′-5′ Construct #2 | RNA | In-house | AAUUAUCACN*NCCCGCAUGCGACA | Figure S5 |

| | | | | |
|---|---|---|---|---|
| 2′-5′ Construct #3 | RNA | In-house | AAUUAUCACN*N*NCCGCAUGCGACA | Figure S5 |
| 2′-5′ Construct #4 | RNA | In-house | AAUUAUCACN*NCN*NGCAUGCGACA | Figure S5 |
| 2′-5′ Construct #5 | RNA | In-house | AAUUAUCAN*NCCN*NGCAUGCGACA | Figure S5 |
| 2′-5′ Construct #6 | RNA | In-house | AAUUAUN*NACCCN*NGCAUGCGACA | Figure S5 |
| RT Test Oligo RC | RNA | IDT | UGUCGCAUGCGGGGGGUGAUAAUU | Figure S5; not truly RC but worked with all constructs; same as NPOM Test Oligo RC |
| CT Mimic and CTB Mimic Primer | RNA | IDT | Alexa488-CUAGUCGCAUGC | Figures 4 and S8 |
| CT Mimic | RNA/DNA | In-house | UCdT(-NPOM)ACCGGCAUGCGACUAG | Figure S8 |
| CT Mimic RC | RNA | IDT | CUAGUCGCAUGCCGGAAGA | Figure S8 |
| CTB Mimic | RNA | In-house | UCdT(-NPOM)CdT(-NPOM)AAAACCGGCAUGCGACUAG | Figures 4 and S8 |
| CTB Mimic RC | RNA | IDT | CUAGUCGCAUGCCGGUUUUAGAGA | Figures 4 and S8 |
| 5′ Handle Block Test | RNA | RNA | GUCGGACUGUAGAACUCUGAAC | Figure S8; not PAGE-purified |
| CT | RNA/DNA | In-house | GUUCAGAGUUCUACAGUCCGACGAUCdT(-NPOM)ACCGGCAUGCGACUAAACGUCGCAUGC | |
| CTB | RNA/DNA | In-house | GUUCAGAGUUCUACAGUCCGACGAUCdT(-NPOM)CdT(-NPOM)AAAACCGGCAUGCGACUAAACGUCGCAUGC | |
| CTEx | RNA/DNA | In-house | GUUCAGAGUUCUACAGUCCGACGAUCdT(-NPOM)CdT(-NPOM)AUACUCCGCAUGCGACUAAACGUCGCAUGCGGAGUA | |
| 6N | RNA/DNA | In-house | GUUCAGAGUUCUACAGUCCGACGAUCdT(-NPOM)CdT(-NPOM)ANNNNNNGCAUGCGACUAAACGUCGCAUGC | |
| 5′ Handle Block | RNA | IDT | GUCGGACUGUAGAACUCUGAA-dideoxyC | |
| RT Handle | DNA | IDT | App-AGATCGGAAGAGCACACGTCT-dideoxyC | |
| RT Primer | DNA | IDT | AGACGTGTGCTCTTCCGATCT | |
| PCR Primer 1 (SR Primer) | DNA | NEB | AATGATACGGCGACCACCGAGATCTACACGTTCAGAGTTCTACAGTCCG-s-A | |
| PCR Primer 2 (Index Primer) | DNA | NEB | CAAGCAGAAGACGGCATACGAGAT(6-base index)GTGACTGGAGTTCAGACGTGTGCTCTTCCGATC-s-T | |

† IDT oligos ordered as RNase-free HPLC-purified unless otherwise noted
‡ RC = Reverse Complement, as competitor during PAGE analysis
⊥ See Material and Methods for details on in-house oligo synthesis and purification
* 2′-5′ linkage
App = riboA 5′-adenylation
dT(-NPOM) = NPOM-caged deoxyT
N = rA, rU, rC or rG
-s- = thiol backbone linkage to inhibit exonucleases

### Accounting of sources of error in NERPE-Seq

It is important to understand sources of error and set a threshold on noise so that output data can be interpreted correctly. Quality filtering aims to minimise reads with errors from sequencing. Additional errors can stem from the protocol, barcode hopping, oligo synthesis (only relevant for some defined-template experiments, see below) and reverse transcription. Our protocol minimises side-products by using rigorous purification steps. We also account for PCR artefacts (Figure S6), and limit barcode hopping by minimising the concentration of residual primers from PCR (Figure S7C) and implementing a barcode read quality filter during Pre-processing. One estimate of barcode hopping in our assay is the false positive rate (read pairs showing any apparent products relative to all read pairs in an experiment) on Pre-processed samples that *should not* exhibit any products because no activated nucleotides were included in the experiment. For CTBC, run with seven other samples, the false positive rate was 0.22% and for 6NC, run with four other samples, it was 0.047%. These are upper bounds on barcode hopping because false positives could also arise from bridge-amplification faults on the sequencing flowcell.

Not all sources of error are directly relevant to interpreting our experimental results. For example, oligonucleotide synthesis errors will not affect our experiments with randomised templates (see below). However, the hairpin constructs all harbour defined sequences (Figure 3) as well as products from primer extension. Furthermore, the Pre-processing compares the known sequences with what is actually measured by sequencing. This enables us to measure sources of error that are of general interest for experiments involving deep sequencing, solid-state oligonucleotide synthesis and reverse transcription. We can estimate the combined sequencing, oligo synthesis and reverse transcription error (under 2´-5´-optimised conditions) by measuring how many reads fail to agree with the defined Fix 1 sequence and are therefore filtered. The average per-base error in Fix 1 across eight samples by this metric is 0.68 ± 0.22%. By comparing three pairs of otherwise orthogonal experiments in which both 2´-5´-optimised and standard reverse transcription conditions were tested, we find that the contribution from the 2´-5´-optimised conditions to the per-base error is 0.24 ± 0.071%. However, we can also measure the combined sequencing, oligo synthesis and reverse transcription error rate *after the reads have passed through all the quality filters during Pre-processing* by looking at samples with defined templates (all of them except 6NC) and measuring deviations from the expected sequences. Here, the per-base error across seven samples is 0.27 ± 0.059%, about half the value measured at the Fix 1 filtering step. We can again measure the contribution from 2´-5´-optimised reverse transcription conditions as above and find a per-base error of only 0.062 ± 0.040%. The lower error rates after Pre-processing suggests error correlation: reads that survive filtering are less likely to harbour errors from sequencing, oligo synthesis and/or reverse transcription. We conclude that under standard reverse transcription conditions, primer extension products with multiple 2´-5´ linkages may be underrepresented (Figure S5), whereas the 2´-5´-optimised conditions bring those sequences along but introduce a small but measurable error factor. We argue that it is generally useful to minimise the potential underrepresentation of any particular product type, but choosing one condition over the other does not complicate the protocol: this variable can simply be toggled during sample preparation, or both conditions can be applied, depending on the questions being asked. Importantly, after Pre-processing is complete, the error contribution from the 2´-5´-optimised reverse transcription conditions is minimal.

To better understand the errors directly related to interpreting results, we can look at primer extension products, which contain errors from sequencing, reverse transcription and primer extension (but not oligo synthesis). We can eliminate the primer extension contribution by tallying physically impossible products: only 2AIrG and 2AIrC were used in the CTB20, CTB40 and CTB60 experiments, but not 2AIrA and 2AIrU. Therefore, any rA or rU products must arise from errors. *By this metric, the experimental per-base error rate from sequencing and reverse transcription across the three experiments is only 0.062 ± 0.13%.* These experiments also enable us to eliminate the possibility that this error arises from nucleotide impurities (for example, contaminating rA in the rG or rC stocks). If impurities did contribute to this error, then we would expect the proportion of apparently impossible errors to increase as overall nucleotide concentration is increased (because then the contaminating nucleotide concentration would also increase) However, the contribution of physically impossible products is independent of nucleotide contribution, registering as 0.070 ± 0.070%, 0.047 ± 0.062% and 0.070 ± 0.084% for 20 mM, 40 mM and 60 mM total nucleotide, respectively.

The contribution of errors from the sequencer is calculated in the Results under "*The 6N template and oligonucleotide synthesiser errors*", where the number of filtered forward reads is compared with the number of subsequently-filtered reverse reads based on the Fix 1 sequence. We find an average per-base sequencing error of only 0.00089%, well below other sources of error and so minor as to be negligible when interpreting NERPE-Seq data.

Taken together, this overview of data noise shows that incorrect base calls in the products will arise from (i) reverse transcription, contributing a fraction of a tenth of a percent error per base, (ii) barcode hopping and bridge-amplification faults, contributing on the order of a tenth of a percent and (iii) sequencing itself, contributing negligibly (see Results). We conclude that interpreting the identity of individual bases in NERPE-Seq results comes attached with an error on the order of a fraction of a percent, which we anticipate will be wholly sufficient for characterising future results.

Our test experiments used constructs with defined templates to enable protocol validation, but some of the presented analyses are frustrated by oligo synthesis errors in a given template. An obvious solution is to simply filter the data for a specific template, and we have included a code module for this. However, NERPE-Seq is designed for randomised templates, which is why the default analysis does not assume any specific template. *Crucially, there is no such category as "template synthesiser error" in the case of a randomised template because no particular sequence is specified.* The only concern is that the overall base distribution will not be truly random, and the analysis accounts for this by calculating normalisation factors for each position.

**Annotation of the NERPE-Seq analysis code**

**Sections**

**A.** General information

**B.** Pre-processing

**C.** Characterisation

### A. General information

The RNA constructs used in these experiments are hairpins that prime NonEnzymatic RNA Primer Extension (NERPE, or PE for short) across an internal template region. Consequently, the template for PE and the product of PE are physically connected and on the same RNA strand. Each sequencing read will contain both template and product sequences, or their reverse complements, enabling us to study which templates yield which products.

**A.1** After passing through the protocol (handle ligation, reverse transcription and PCR), the dsDNA to be sequenced looks essentially like this:

```
                 10        20        30        40        50
                  |         |         |         |         |
5'    AATGATACGGCGACCACCGAGATCTACACGTTCAGAGTTCTACAGTCCGA
3'    TTACTATGCCGCTGGTGGCTCTAGATGTGCAAGTCTCAAGATGTCAGGCT

                 60        70        80        90        100
                  |         |         |         |         |
      CGATCTCTANNNNNNGCATGCGACTAAACGTCGCATGCNNNNNNNAGATC
      GCTAGAGATNNNNNNCGTACGCTGATTTGCAGCGTACGNNNNNNNTCTAG
           Prefix    Fix 1                 Product
               Template

                 110       120       130       140       150
                  |         |         |         |         |
      GGAAGAGCACACGTCTGAACTCCAGTCACNNNNNNNATCTCGTATGCCGT
      CCTTCTCGTGTGCAGACTTGAGGTCAGTGNNNNNNNTAGAGCATACGGCA
      Fix 2                         Barcode

                 160
                  |
      CTTCTGCTTG 3'
      GAAGACGAAC 5'
```

**A.1.1 Variable 1 Among Constructs: Template.** The template is of *fixed length* for a given RNA construct. However, the number and identity of bases in the template varies across constructs and is usually reflected in the construct name (for example, 6N uses a construct with 6 randomised bases as a template).

**A.1.2 Variable 2 Among Constructs: Prefix.** The Prefix motif contains two T residues that in the parent hairpin construct harbour a physical block to primer extension (the NPOM caged bases).

**A.1.3 Variable 3 Among Constructs: Maximum Allowable Product Length.** (See above, A.1.2.) *Primer extension cannot extend beyond the first T that is encountered* (see Figure S3). However, formally, primer extension can potentially extend up to and including the A residue preceding the T, immediately adjacent to the last base of the template. *The maximum possible product length is therefore equivalent to the length of the template + 1.* The product is of variable length within an experiment (because it is a function of primer extension efficiency) and also across constructs (because it depends on the nature of the template); its length is between 0 and [(length of template) + 1].

**A.2 Sequencing Overview:** The PCR product shown in **A.1** is optimised for paired-end Illumina® sequencing. For documentation of primer sequences see NEBNext® Multiplex Small RNA Library Prep Kit for Illumina® from New England Biolabs®[1]. NERPE-Seq assumes that sequencing files are provided debarcoded and with an index file, so that each sample has three FASTQ files representing the index, the forward and the reverse reads. These files must also be sorted, so the Nth read in one file corresponds to the Nth read in the other files. This is checked during the analysis process to prevent the processing non-sorted files.

**A.2.1 Forward v. Reverse Reads:** Understanding the relationship between the forward and reverse reads and the parent sequences under study is very important. If this is ambiguously defined then the analysis could inadvertently identify patterns in reverse complement (antisense) space rather than in sense space.

**A.2.2** Because the parent hairpin is a single strand of RNA and includes both the sense template and the sense product (which should nominally be complementary to each other), one of the reads necessarily contains both the sense template and the sense product. This happens to be R1. R2 contains the antisense of both. *Therefore R1 represents the parent sequence(s) we are ultimately interested in. R2 is the reverse complement of the parent sequence(s).*

[1] https://www.neb.com/-/media/nebus/files/manuals/manuale7300_e7330_e7560_e7580.pdf?la=en&rev=1924b6d968114d8290cb772f3d85f13c&hash=567FB22B3BBBB307DE905A107261B2ADBEDAF8DA

**Example R1**

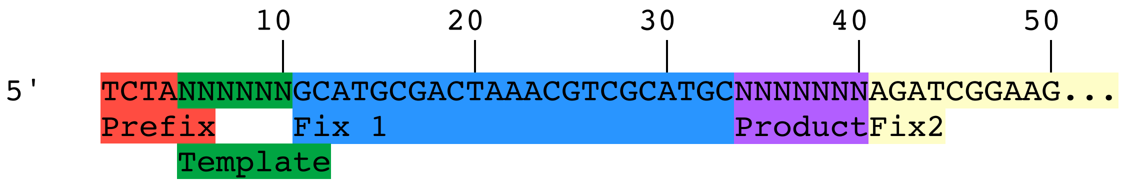


**Example R2**

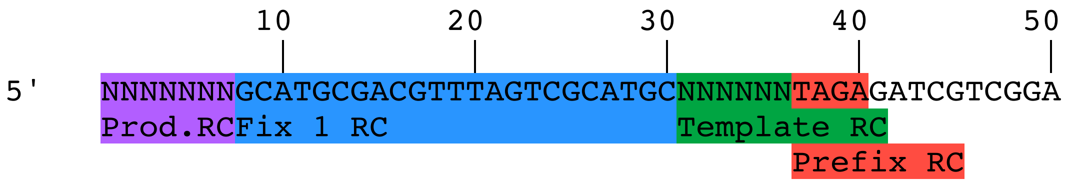


**A.2.3** Reality check: R1 should begin with the sense Prefix sequence.

**A.2.4** Reality check: R2 should begin with the antisense Product sequence.

**A.2.5** Because the parent sequence is a hairpin, the Product and Template "point away" from each other in R1 space. (That is, the hairpin is one continuous sequence, 5´-to-3´, but it folds back onto itself and primer extension happens in the 5´-to-3´ direction on an oppositely-oriented template.) This is critical for correct template-product alignment. *The 3´-most base of the template must be aligned flush with the 5´-most base of the product*.

**A.3 Software overview**

**A.3.1 Software Required.** The NERPE-Seq software package is implemented in MATLAB. It has been tested in MATLAB versions 2019b and 2017a and is likely backward compatible for some earlier versions. The *Bioinformatics*, *Distributed Computing* and *Statistics* toolboxes are also required.

**A.3.2 Analysis.** Each pair of three files (index, forward and reverse) first undergoes Pre-processing (section B) prior to Characterisation (section C). To facilitate processing of a set of barcoded samples, the *ProcessSamples.m* function can be used.

This function utilizes a Microsoft Excel spreadsheet as an input, with one row of headers and one additional row per sample. For each sample one defines the sequencing files, the output directory, the length of the template region, the sequence identity of the Prefix and, as discussed below, a control condition to use for normalisation, and the template normalisation factor.

To facilitate rapid random access to sequencing reads, which is later required for parallel processing of blocks of sequencing reads during Pre-processing, the FASTQ input files are initially indexed into blocks using a custom function *fastqmap()*. Next, each set of input files is analysed using *preprocess()* and *characterize()*. We use *read block* to describe a block of sequencing reads processed at one time.

**B. Pre-processing**

The goal of this section is to generate datasets for downstream analysis and is implemented by the *preprocess()* function (preprocess.m).

The target data for downstream analysis consists of individual sets (compiled in one FASTA-like file per experiment/barcode) that look like this:

> header (ID#: $N^{th}$ read from sequencing file)

5' NNNN*****
3' NNNNNNA

Here, the green sequence is the reverse (not reverse complement) of the template in the forward read (see A.2.5).

Carrying over sequencing quality data is unnecessary because DNA sequencing errors are uncorrelated with errors in primer extension. That is, an error in primer extension is expected to show up in the sequencing data as a mismatch between template and product, but there is no reason to posit that it is therefore likelier to be incorrectly sequenced.

As in **A.1**, purple denotes product and green denotes template. Note the relationship between what this data looks like and the points above. These are sense sequences (we obviously expect them to be reverse complements *of each other*, but they are both sense in that they are represented by R1, and therefore are the sequences that were present in the initial RNA construct). The product runs 5´-to-3´ and the template is correctly oriented with respect to the product. The last base in the templating region is the A of the Prefix that precedes the blocking T. The asterisks represent unextended bases in this representative example.

In the following steps, each read pair must pass a series of sequential checks in order to be considered valid for Characterisation. Any read pair failing a check is discarded and not analysed further.

**B.1.1** <u>Read numbers match.</u> The number of reads in R1 and R2 (R1/2), and the index read file must be identical.

**B.1.2** <u>Read blocks.</u> Next, reads are analysed in *read blocks* using parallel processing of one *read block* per thread, with a default of 10,000 reads per *read block*. Once all reads in a *read block* are pre-processed, a *read block* FASTA-like file is saved, and block summary statistics are generated. When all blocks have been analysed, a summary of all blocks is generated and saved, the intermediate MAT files are archived to .zip format and the individual FASTA-like *read block* files are concatenated.

**B.2.1** <u>Header match.</u> Within each thread, processing is serial, and each read in the *read block* undergoes Pre-processing using a helper function *preprocess_read()*, starting with checking explicitly that the file headers match. This step and the Pre-processing steps below occur in this helper function (preprocess_read.m).

**B.2.2** <u>Index quality filter.</u> Converts quality-scores of index reads to error probabilities (which can be averaged). Discards all index reads with average error probabilities higher than that equivalent to q = 26. The equivalent Phred quality score is computed using the *qpbar()* function (qpbar.m).

**B.2.3** R1 quality filter. Discards all R1 reads with average error probabilities higher than that equivalent to q = 30.

**B.2.4** R2 quality filter. Discards all R2 reads with average error probabilities higher than that equivalent to q = 30.

**B.3** Use Fix 2 as a quality filter on R1. Demands that R1 contain a *single* perfect match to the Fix 2 sequence. (R2 does not contain Fix 2.)

**B.4** Trim R1. Trims R1 where Fix 2 starts (that is, cuts Fix 2 and downstream out).

**Example R1**

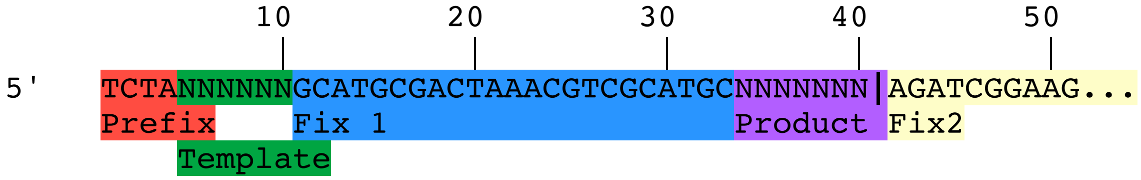


**B.5** Use Fix1 as a quality filter. Demands that both R1 and R2 contain a perfect match to Fix 1.

**B.6** Confirm R1 template (B.6.1) and Prefix (B.6.2). The template and Prefix are of defined length in each construct, but the template can be a randomised sequence. Therefore, we cannot use sequence to identify it, and it must be located with reference to Fix 1.

This step:
- Defines the template by indexing the appropriate number of bases (as defined for the given construct) relative to Fix 1. If doing so walks off the read, it is discarded (means something is wrong with the Prefix and/or template length).

- Defines the Prefix by indexing the appropriate number of bases (as defined for the given construct) relative to template. If doing so walks off the read, it is discarded.
- Confirms Prefix identity. Note that this step automatically also eliminates reads with templates that are an incorrect length (shorter or longer). A template with an incorrect length will cause the Prefix to deviate from the actual, known Prefix sequence/length and such reads will therefore be eliminated.
- Checks that Prefix location in R1 starts at position 1 (B.6.3).

**B.7** Confirms R2 template (B.7.1) and Prefix (B.7.2), as in step B.6, and confirms that a Fix 3 match is found to the right of Prefix in R2 (B.7.3).

**B.8** <u>Trims R1/2</u> in the Prefix such that only the templating A (in R2 this appears as a T) is kept. This is the base immediately adjacent to the template. Everything to the left is trimmed.

**Example R1**

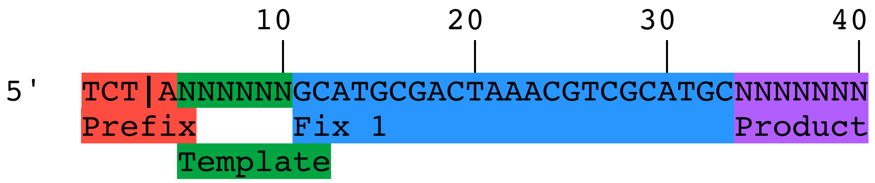


**Example R2**

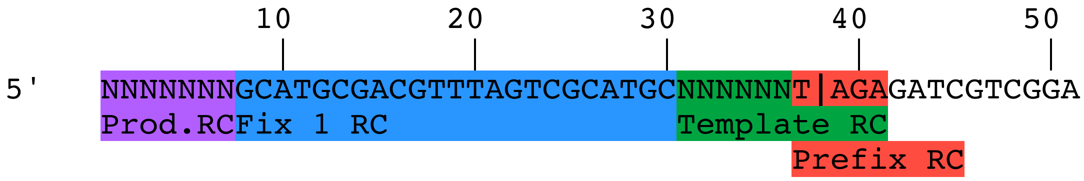


This trim is not introduced sooner because the Prefix sequence is very short—just 4 bases—and is difficult to identify based on sequence alone.

**B.9** Product identification. In R1, the product is between Fix 1 and the end of the read (it has already been trimmed of Fix 2). In R2, the product is between the start of the read and fix 1.

**B.10** Product length filter. See A.1.3: the product length cannot be longer than what is physically allowed. Removing any violating sequences limits potential false positives. This step also demands that product length in R1/2 be ≤ [(length of template) + 1].

**B.11** Product equivalent length filter. Demands that the product length in R1 equals the product length in R2.

**B.12** Final sequence comparison: product and template quality filters. Compares R1 and R2 *to each other* across their entire lengths, and demands the sequences be perfectly complementary. This simultaneously confirms correct product and template complementarity in the reads.

**B.13** Final processing to yield analysable data and conversion to RNA space. We have used both R1 and R2 throughout to parse and quality-control the sequences, but at this point we no longer need both reads. *This step extracts the template (and the templating "A" from Prefix) and product from R1*. The rest of the sequences and most of the metadata are no longer needed. For ease of interpretation all "T"s are converted into "U"s.

**B.14** Base/position naming system. For the ease and convenience of representing and processing data, the first base in the template (3´) is denoted as "T1", and the base that is opposite T1 as "P1"' ("product one"). T2 and P2, and so on. This means numbering on the product proceeds from 5´-to-3´ and for the template from 3´-to-5´.

**C. Characterisation**

Characterisation provides general metrics about primer extension in a given experiment. Note that the following requires the Pre-processed output of section B. Characterisation is implemented in the *characterize()* function (characterize.m).

**C.1** Extension events. This is a proxy for primer extension efficiency.

**C.1.1** Calculates the raw number of unextended, +1 extended, +2 extended, *etc*. products. This is equivalent to the data provided by PAGE analysis.

**C.1.2** Lists C.1.1 as frequencies relative to the total number of products.

**C.2** Template Composition. Understanding the template makeup enables us to interpret the product makeup. (Biases in the product makeup may stem from biases in template makeup, which is a function of solid-state RNA synthesis.) Also, importantly, differences in the template makeup of a negative control and the template makeup of experimental samples may reflect biases in RT activity on 2´-5´ linkages.

**C.2.1** Lists the raw position-dependent base composition of the template.

**C.2.2** Lists the position-dependent base composition of the template (C.2.1) as frequencies.

**C.2.3** Compares the position-dependent base composition of the template to that of any specified control template (if specified) by taking the ratios of the bases at each position. The null hypothesis is that the ratios will equal 1, whereas any deviation from unity could point to a sequence-dependent 2´-5´ linkage signature.

NOTE: Certain template sequences may preferentially yield products that are a problem for one or another enzyme during sample preparation. Therefore, the set of template sequences that survives sample preparation for an experimental sample may be different from the set of template sequences that survives in a control sample (where there cannot be any biases in enzyme activity due to primer extension because primer extension did not happen). That is, the template makeup as measured for the control sample is more correctly representative of the true randomness of templates available to primer extension during the experimental reaction.

**C.2.4** Uses C.2.2 to extract the *normalisation factors* for each base at each position; these factors represent template deviation from an ideal, truly random makeup. The ideal frequency for each base at each position will usually be 0.25, but this may be varied experimentally. Note that all downstream normalisations are performed using the control template.

In the input Excel file (see **A.3.2**), these are specified in the 3´-to-5´ direction so the base sequence or the base probability distribution is specified for product position 1, 2, 3, … in the 5´-to-3´ direction of the product. This means that the normalisation factor is the reverse (not reverse complement) of the template sequence synthesized.

**C.3** Product grouping.

**C.3.1** Group complementary, mismatch-containing and unextended blocks.
- Blocks with perfectly complementary products:templates = "**Complementary Set**".
- Blocks with one or more mismatches = **"Mismatch Set".**
- Blocks with no product = **"Unextended Set".**

**C.3.2** Counts how many blocks in each group. (Reality check: these three groups account for all possible types and should add up to the total number of products in a given experiment.)

**C.4** Product features in the **Complementary Set** and **Unextended Set**.

**C.4.1 Complementary Set and Unextended Set.** Computes the raw position-dependent base composition of the products, *including nulls*.

**C.4.2 Complementary Set.** Normalises the raw position-dependent base composition of the product (C.4.1) to the template using the normalisation factors (C.2.4), *excluding nulls*. In this case we must permute the rows of the template normalisation factor to account for the fact that we are looking at the product and not the template *(i.e.*: complementary bases).

**C.4.3 Complementary Set.** Computes the normalised position-dependent base composition of the product (C.4.2) as frequencies.

**C.4.4 Complementary Set.** Normalises the raw position-dependent base composition of the product (C.4.1) to the *control template* using the *control template normalisation factors* (C.2.4).

See also NOTE at C.2.3.

**C.4.5 Complementary Set.** Computes the *control-template-normalised* position-dependent base composition of the product (C.4.4) as frequencies.

**C.5** Sequence-dependence of nulls (primer extension termination/stalling) in the **Complementary Set and Unextended Set.** This analysis allows us to answer the question: Do any terminal product bases or templating bases tend to stop primer extension?

**C.5.1 Complementary Set.** Computes the raw position-dependent base distribution of all terminal product bases (*i.e.*: the base distribution of the final base of all products at each position).

**C.5.2 Complementary Set.** Normalises the raw position-dependent base distribution of all terminal product bases (C.5.1) to the template using the normalisation factors (C.2.4).

**C.5.3 Complementary Set**. Computes the normalised position-dependent base distribution of all terminal product bases (C.5.2) as frequencies.

**C.5.4 Complementary Set and Unextended Set**. Computes the raw position-dependent base distribution of all ***templating bases*** one base downstream of all terminal products (which, formally, in this case, include all unextended products).

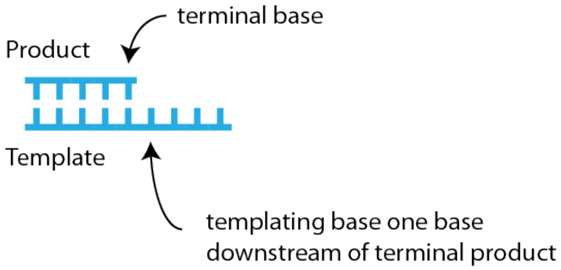


**C.5.5 Complementary Set and Unextended Set.** Normalises the raw position-dependent base distribution of all templating bases one base downstream of all terminal products (C.5.4) to the template using the normalisation factors (C.2.4).

**C.5.6 Complementary Set and Unextended Set.** Computes the normalised position-dependent base distribution of all templating bases one base downstream of all terminal products (C.5.5) as frequencies.

**C.5.7 Complementary Set.** Computes the frequency of each type of dimer in the template as a function of incorporation position i, where each dimer is the template sequence at positions [i, i+1]. Computes both position-dependent count and frequency.

**C.5.8.** Computes the number of template dimers (entire set) as a function of incorporation position i, where each dimer is the template sequence at positions [i i+1]. Computes both position-dependent count and frequency.

**C.6 Mismatch Set**. Mismatches. What are the properties of products among mismatches?

**C.6.1 Mismatch Set**. Computes how many products, relative to the total of all extended products (Complementary Set + Mismatch Set), have 1, 2, 3, *etc*. mismatches.

**C.6.2 Mismatch Set**. Computes the position-dependent distribution of terminal mismatches.

**C.6.3 Mismatch Set**. Computes the position-dependent distribution of a mismatch followed by a mismatch.

**C.6.4 Mismatch Set**. Computes the position-dependent distribution of a mismatch followed by a correctly paired base.

**C.6.5 Mismatch Set**. Sequence space of Mismatches. What is the position-dependent prevalence of each possible mismatch? This step defines each possibility as a bin at each position, and generates counts in each bin. To normalise, it applies normalisation factors across the entire contents of each base-defined set of bins (*e.g.*: in position 1, the "A bin" will contain AA, AG, and AC in some proportion of raw counts of each—normalises all three bins to the appropriate position 1 normalisation factor).

**C.7 Trimer Frequencies Visualisation.** A custom function seqspace_cube() generates a visualisation showing trimer (stretches of three bases) frequencies at each position, where the volume of each sphere is proportional to the frequency. For a template length of 6, this would include position k = {1, 2, 3, 4}, up to template length - 2. Below is an example plot for a randomised template at k = 1 from the 6NC.

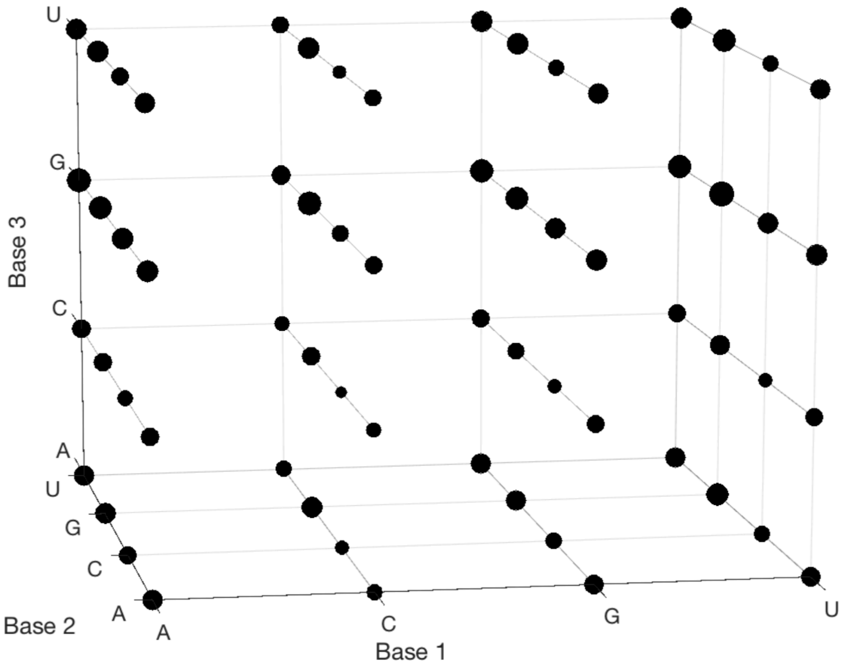


**C.7.1** Generates sequence space cube (trimer) plots and data (all sets, including the templates).

**C.7.2** Generates sequence space cube (trimer) plots and data for Complementary set.

**C.8 Transition Probabilities.** Calculates the position-dependent base transition probabilities at each position. Generates transition map plots. At any given position, the next base can be A, C, G, U or null (unextended, for product only). Using a frequentist interpretation of probability, the transition probability is calculated as the number of transitions observed normalised by the number of observations at a given position. This is done in the helper function *transition_map()*. Counts and then frequencies are separately computed and visualised for the product (with and without nulls) and template.

### SUPPLEMENTARY REFERENCES


1. Tessier, D.C., Brousseau, R. and Vernet, T. (1986) Ligation of Single-stranded Oligodeoxyribonucleotides By T4 RNA Ligase. *Analytical Biochemistry*, **158**, 171-178.
2. Lorsch, J.R., Bartel, D.P. and Szostak, J.W. (1995) Reverse-transcriptase Reads Through a 2'-5'-Linkage and a 2'-thiophosphate in a Template. *Nucleic Acids Res.*, **23**, 2811-2814.
